# Disturbance-based management of ecosystem services and disservices in partial nitritation anammox biofilms

**DOI:** 10.1101/2021.07.05.451122

**Authors:** Carolina Suarez, Christopher J. Sedlacek, David J. I. Gustavsson, Alexander Eiler, Oskar Modin, Malte Hermansson, Frank Persson

## Abstract

The resistance and resilience provided by functional redundancy, a common feature of microbial communities, is not always advantageous. An example is nitrite oxidation in partial nitritation-anammox (PNA) reactors during wastewater treatment, where suppression of nitrite oxidizers like *Nitrospira* is sought. In these ecosystems, biofilms provide microhabitats with oxygen gradients, allowing the coexistence aerobic and anaerobic bacteria. We designed a disturbance experiment where PNA biofilms treating water from a high rate activated sludge process removing organic matter (mainstream wastewater), were constantly or intermittently exposed to the effluent of anaerobic sewage sludge digestion dewatering (sidestream wastewater), which has been proposed to inhibit nitrite oxidizers. With increasing sidestream exposure we observed decreased abundance, alpha-diversity, functional versatility, and hence functional redundancy, among *Nitrospira* in the PNA biofilms, while the opposite patterns were observed for anammox bacteria within *Brocadia*. At the same time, species turnover was observed for the aerobic ammonia-oxidizing *Nitrosomonas* populations. The different exposure regimens were associated with metagenomic assembled genomes of *Nitrosomonas, Nitrospira*, and *Brocadia*, encoding genes related to N-cycling, substrate usage, and osmotic stress response, possibly explaining the three different patterns by niche differentiation. These findings imply that disturbances can be used to manage the functional redundancy of biofilm microbiomes in a desirable direction, which should be considered when designing operational strategies for wastewater treatment.

## INTRODUCTION

The insurance hypothesis states that biodiversity can protect against the loss of ecosystem services even when ecosystems are exposed to disturbances ^1^. This is because functional redundancy, a direct result of biodiversity, is predicted to provide resilience and resistance against environmental change ^2,3^. Furthermore, biodiversity is linked to multifunctionality and the capability of a community or population to provide a multitude of ecosystem services ^4-6^. In highly diverse microbial systems such as soils, sediments and wastewater treatment bioreactors, a high degree of multifunctionality and functional redundancy should consequently lead to high resilience and a resistant supply of ecosystem services. These are preferred features for many ecosystems, protecting them from anthropogenically induced changes ^7,8^. However, in engineered systems certain ecosystem services are unwanted (so called disservices) ^9,10^. Functional redundancy of disservices within an engineered system can hinder the tuning of these systems towards more desirable operational function.

Wastewater treatment plants (WWTPs) are engineered ecosystems providing fundamental ecosystem services for the built environment ^11^. Enhanced biological nitrogen removal is used to reduce the amount of reactive nitrogen in the effluent of WWTPs, which otherwise would lead to reduced water quality and eutrophication in recipient waterways ^12^. Nitrogen removal is commonly achieved by maintaining two ecosystem services: nitrification and denitrification. However, the recent focus on energy neutrality at WWTPs has promoted processes based on nitritation. Here the influent ammonium is oxidized to nitrite by ammonia-oxidizing bacteria (AOB), but the further oxidation of nitrite to nitrate by nitrite-oxidizing bacteria (NOB) is undesired ^13^ and hence an ecosystem disservice. Successful suppression of NOB leads to the accumulation of nitrite, which can be reduced to nitrogen gas in denitrification systems ^14^ or by anaerobic ammonia-oxidizing (anammox) bacteria in partial nitritation-anammox (PNA) bioreactors ^15^. Because of their slow growth rate, biofilm systems are commonly used to maintain anammox bacteria and AOB in WWTP bioreactors ^16^. Moreover, biofilms provide oxygen gradients allowing the coexistence of these bacterial groups in the same biofilm ^17^.

Nitrification is performed by ammonia- and nitrite oxidizers belonging to multiple genera, where *Nitrosomonas, Nitrospira* and *Nitrotoga* are the most common in WWTPs. Furthermore, multiple subpopulations are often reported to coexist for the AOB *Nitrosomonas* ^18,19^ and the NOB *Nitrospira* ^20^. This microdiversity within nitrifier populations might represent ecotypes, closely related populations that can occupy different ecological niches ^21^. For example, closely related *Nitrosomonas* and *Nitrospira* populations are known to differ in their preferences for substrate concentration ^22,23 24^ and temperature ^23,25^, as well as their ability to use alternative energy sources such as urea, cyanate, formate, and hydrogen ^24,26-28^. In addition, some *Nitrospira* are nitrite oxidizers, while other closely related *Nitrospira* are capable of complete ammonia oxidation to nitrate ^29-32^. In the case of anammox bacteria, microdiversity within wastewater treatment microbiomes is less documented but has been demonstrated in marine ecosystems ^33,34^. It is also evident that ecotypes within *Brocadia*, common in anammox wastewater treatment reactors, differ in their substrate preferences and responses to reactor operation ^35,36^. As opposed to the autotrophic processes of nitrification and anammox, denitrification is a process carried out by a large number of taxa, distributed across multiple phyla, with considerable metabolic versatility ^37^.

In PNA bioreactors treating municipal wastewater (“mainstream water”), modelling approaches and experimental results have shown that it is difficult to inhibit NOB while simultaneously maintaining high activities of AOB and anammox bacteria, due to the cold water having relatively low ammonium and free ammonia (NH_3_ and FA) concentrations ^16,38^. Exposing mainstream communities to the sludge liquor from sewage sludge anaerobic digestion (“sidestream water”), which is warm and ammonium rich, has previously been proposed as a method to inhibit NOB in wastewater treatment reactors ^39,40^. The inhibition could occur due to multiple mechanisms, such as competition for nitrite with anammox bacteria, increased competition for oxygen with AOB, or the presence of a higher FA concentration ^41-43^. Therefore, a disturbance management with sidestream exposure may reduce the biodiversity and functional redundancy of NOB in PNA bioreactors.

However, for a disturbance management with sidestream exposure approach to be viable in PNA bioreactors, the functional diversity of AOB and anammox bacteria must not be negatively affected. Although sidestream exposure is expected to shift the composition of NOB, AOB, and anammox bacteria, how it will affect their abundance, biodiversity, and functional redundancy is unknown. Since detailed information on such ecological fundamentals is scarce, NOB inhibition attempts in PNA bioreactors are largely based on trial and error.

At the Sjölunda WWTP (Malmö, Sweden), mainstream and sidestream pilot PNA reactors harboured biofilm communities with *Nitrospira, Nitrosomonas* and *Brocadia* populations, which have been previously characterized to contain various degrees of microdiversity ^44^. In this study, we designed a disturbance experiment where mainstream PNA biofilms were intermittently or constantly exposed to sidestream wastewater (Figure 1). Our aim was to investigate whether such a disturbance regimen could be utilised to shift the microbial community of biofilms towards a desired state. More specifically, we describe the effect of a gradient of sidestream exposure on the abundance, microdiversity, and functional diversity of NOB, AOB, anammox bacteria and denitrifiers.

**Figure 1.**
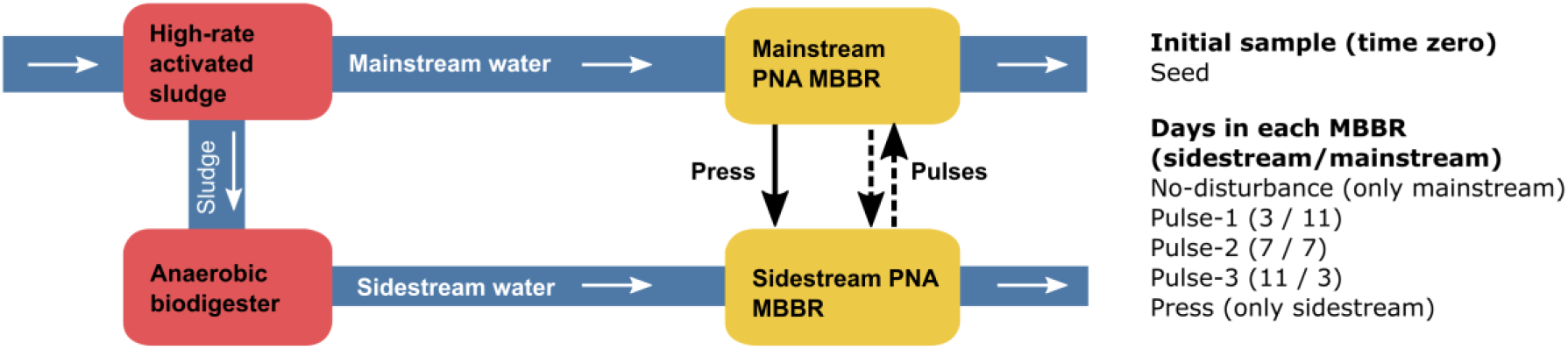
Scheme of the MBBRs at the Sjölunda WWTP and the experimental design. White arrows indicate the water flow. The black arrows indicate transfer of biofilm carriers between the PNA bioreactors (yellow). Biofilm carriers were either kept in the mainstream PNA MBBR (No-disturbance), moved to the sidestream PNA MBBR (Press), or moved between the mainstream and the sidestream in two-week periods according to the time schedule (Pulse-1 to Pulse-3). Initial Seed samples, at time zero were also analysed. The two-week rotation periods were repeated 4 times over the course of 58 days.

We hypothesize that although complete suppression of *Nitrospira* is unlikely, certain populations or ecotypes of *Nitrospira* will be sensitive to sidestream wastewater. If so, the microdiversity of *Nitrospira* and thereby their functional redundancy will decrease. Sidestream exposure is also expected to shift the composition of *Nitrosomona*s and *Brocadia* communities, but how their functional redundancy will be affected is unknown. We also hypothesise that the denitrifying microbiome, which is dispersed across multiple phyla, will be less affected by these disturbances.

## RESULTS AND DISCUSSION

Long-term and short-term disturbances are known as press- and pulse disturbance, respectively ^45^. In this study, we exposed biofilm carriers from a mainstream PNA moving bed biofilm bioreactor (MBBR) to sidestream wastewater in another MBBR at varying time intervals over the course of 58 days (Pulse disturbances 1, 2, and 3). In addition, biofilms carriers were moved from the mainstream to the sidestream MBBR for the entire 58-day period (Press disturbance). Lastly, a set of biofilm carriers were kept in the mainstream MMBR for the entire 58-day period as a control (No-disturbance) (Figure 1).

### Rare taxa were particularly sensitive to sidestream wastewater

In PNA biofilms, nitrifiers are restricted to a thin oxic layer at the surface of the biofilm, and hence their relative abundance in the biofilm is low ^44,46^; therefore, beta-diversity metrics, sensitive to rare taxa, are well-suited to elucidate their distribution patterns. For metrics based on Hill-numbers, this sensitivity can be controlled with the q parameter (diversity order). When q=0, which is a presence-absence metric, the index is insensitive to the relative abundances of taxa (equivalent to the Sørensen index) while for increasing q, the index is more sensitive to abundant taxa^47,48^.

The PNA mainstream biofilm community appears to be sensitive to sidestream water, as we observed differences between treatments for both rare and abundant taxa in the 16S dataset (PERMANOVA q=0, 1 and 2; p<0.01). Excluding the Seed samples at time zero, there is a beta-diversity gradient between no-disturbance and press-disturbance, corresponding to the extent of sidestream exposure (Figure 2A). Between-group beta-diversity was the highest at q=0 and was also higher than within group dissimilarity at low q values (Figure 2B), implying that rare taxa were more sensitive to sidestream exposure than abundant taxa. Similar beta-diversity patterns were seen in the metagenome dataset (Figure 2C, 2D), with differences for rare taxa (PERMANOVA q=0 and q=1; p<0.01), but not the abundant ones (PERMANOVA q=2; p=0.16). Rare taxa have been shown to have important contributions to population dynamics and beta-diversity in microbial communities ^49,50^, as also shown in this study.

**Figure 2.**
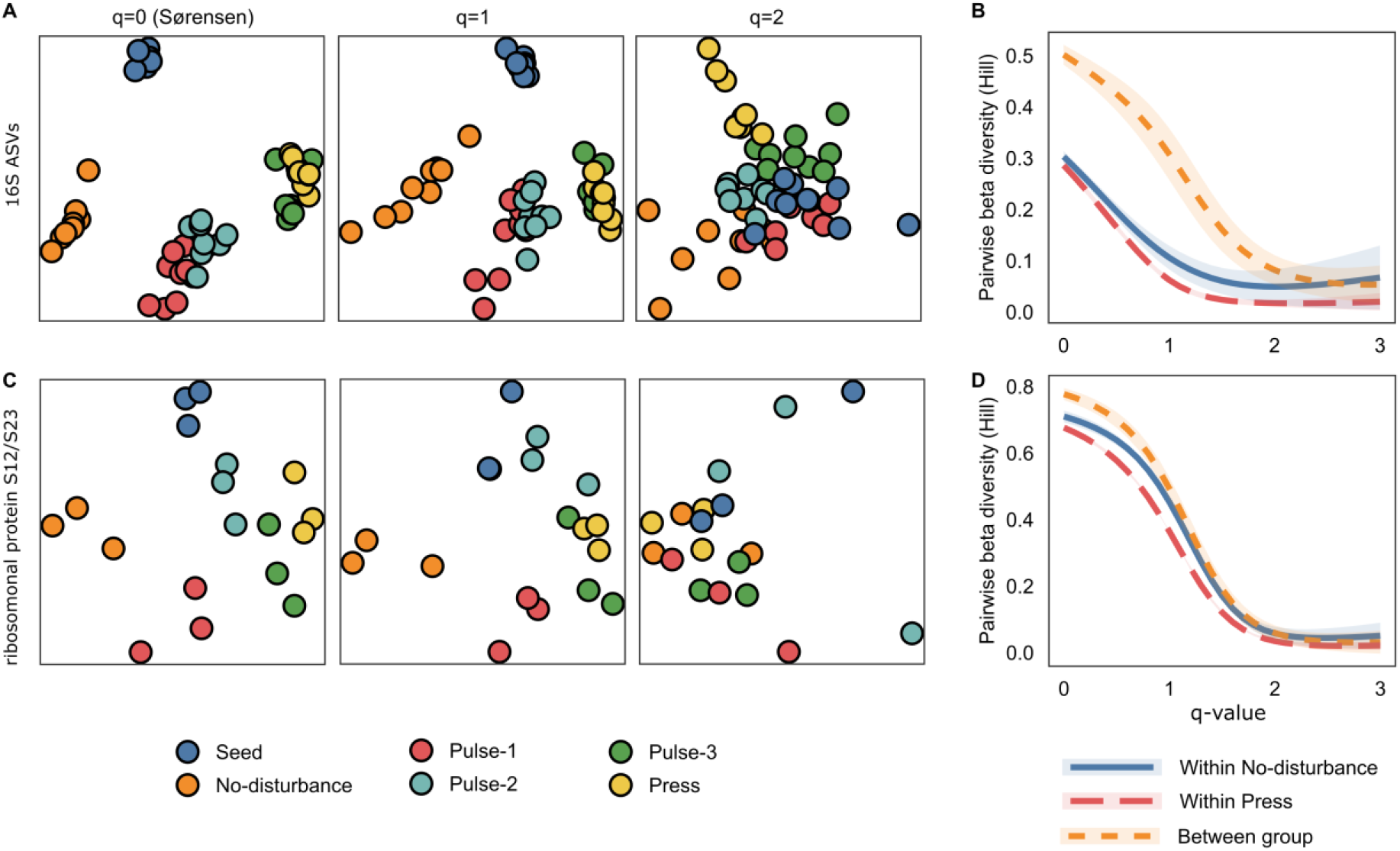
Beta-diversity of 16Sr RNA ASVs (A, B) and of the metagenomics dataset using the ribosomal protein S12/S23 (C, D), based on Hill numbers. A, C: PCoA when q=0 (i.e. the Sørensen index), q=1 or q=2; each circle corresponds to a biofilm carrier. B, D: Within group and between group beta-diversity for No-disturbance and Press-disturbance at different q-values; the lines show the average pairwise beta-diversity and the shaded areas show 95% confidence intervals.

The high within-group beta-diversity observed with both methods could be the result of stochastic assembly of the microbial communities on the replicate biofilm carriers, as also suggested for other biofilms and bioreactors ^17,51^. However, the within-group beta-diversity may also be influenced by methodological issues such as PCR errors for the 16S dataset, sequencing errors, or overestimation of beta-diversity due to undersampling ^52^. For the metagenome dataset, where under-sampling may play a larger role than in the 16S dataset (Figure S1), this could help in explaining why beta-diversity was higher at low q-values and the differences were smaller between groups compared to within groups.

### Response in alpha diversity and abundance to sidestream exposure varies between taxa

We used singleM to determine the abundance and alpha-diversity of nitrifiers in the metagenome dataset. The relative abundance of *Nitrospiraceae* in the biofilms decreased with sidestream exposure, although complete suppression was not achieved (Figure 3A). This lower abundance was associated with lower alpha-diversity (Figure 3B). As an alternative to the singleM-based assessment of species diversity, we also used Kaiju to assign taxonomy to individual genes in the metagenome assembly, and thus estimate a rough distribution of those genes across samples. From this analysis, it was clear that the number of *Nitrospira* genes also decreased with increasing sidestream exposure (Figure 3C).

**Figure 3.**
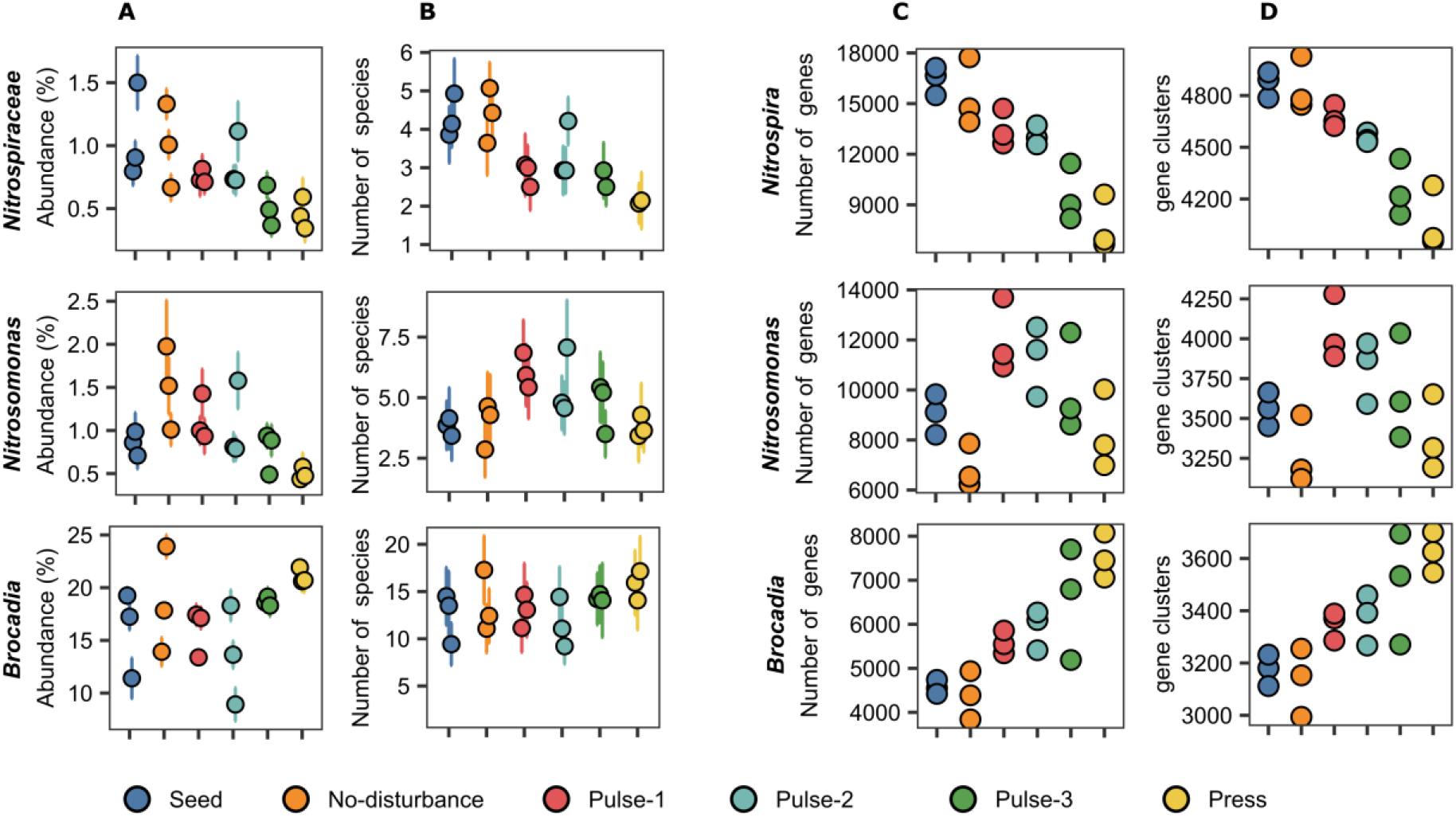
Abundance and richness of *Nitrospira* (top), *Nitrosomonas* (middle), and *Brocadia* (bottom) in the metagenome dataset. **A:** Relative abundances. **B:** The number of species. **C:** The number of genes. **D:** Number of pangenome gene clusters. **A-B:** singleM data; circles show average values of 14 marker genes, error bars indicate 95% confidence intervals. **C-D**: Kaiju data. Note that Kaiju uses NCBI taxonomy, while SingleM uses GTDB taxonomy; in the latter the genus *Nitrospira* is classified as the family *Nitrospiraceae*.

In contrast to the *Nitrospira*, for the AOB *Nitrosomonas*, higher numbers of species (alpha-diversity) and genes (functional diversity) occurred at intermittent exposure to sidestream water (Pulse 1-3; Figure 3), akin to predictions by the intermediate disturbance hypothesis ^53,54^. Among the anammox bacteria (*Brocadia*) an average of ∼15 genotypes were detected across all treatments, but the number of genes assigned to the genus *Brocadia* steadily increased with increasing sidestream exposure (Figure 3). Thus, positive (*Brocadia*), intermediate (*Nitrosomonas*) and negative (*Nitrospira/Nitrospiraceae*) effects on intra-genera diversity were clearly observed in response to the varying levels of disturbance.

The pangenome, coined to refer to all the genes that can be found in a clade including the core and all accessory genes ^55,56^, is assumed to be responsible for niche differentiation and allows the discrimination of different ecotypes, defined as populations of cells adapted to a given ecological niche. We grouped all genes taxonomically classified as *Nitrospira* into gene clusters, which could be considered as the *Nitrospira* pangenome. We show that the number of *Nitrospira* gene clusters also has a downward trend (Figure 3D), suggesting a decrease in functional potential with increased sidestream exposure. This reduction in functional redundancy within *Nitrospira* is beneficial when managing a disservice, as is the case of nitrite oxidation in PNA processes. The opposite pattern occurs for *Brocadia*, leading to increased diversity (Figure 3D), and thus enhanced redundancy in the functional group of anammox bacteria.

The existence of NOB, AOB, and anammox ecotypes with overlapping N-cycling functions (but adapted to different niches) likely confers stability to microbial ecosystems ^57^, resulting in long-lasting persistence of these functional groups in PNA biofilms. Still, even though these functional groups may appear as highly redundant, only a few members may actively be performing specific steps of the N-cycle at a given time. Some members could be exhibiting alternative modes of metabolism ^26,27^, while others may simply be inactive. The metabolic functions performed by a given population generally depends on environmental conditions as well as on the presence and activity of other community members ^58^.

At the Sjölunda WWTP, relative nitrate production was higher in the mainstream PNA MBBR than in the sidestream PNA MBBR ^44^. In addition, other studies have shown that for mainstream communities, exposure to sidestream water leads to nitrite accumulation due to the inhibition of NOB ^39,40^. Together, these data highlight how sidestream wastewater disturbance management regimens can be used to inhibit NOB populations in PNA biofilms and preserve AOB and anammox.

### Distinct *Nitrosomonas* populations were present in the sidestream and mainstream

The disturbances resulted in differences in richness (nestedness) among the nitrifying- and anammox communities (above). However, differences between communities (beta-diversity) can also arise because of species turnover, the latter occurring when one species is replaced by another ^59^. When comparing No-disturbance with Press-disturbance for *Nitrosomonas* genes, the beta-diversity observed was largely due to turnover (Figure 4), as also seen in Figure S2. Notably, this was not the case for the *Brocadia* or *Nitrospira* species (Figure 4). Thus, distinct *Nitrosomonas* populations seem to be present in the biofilms exposed to sidestream and mainstream wastewater. Similar results were observed in the 16S dataset, were a single *Nitrosomonas* ASV was dominant in the mainstream wastewater and its abundance was lower in all samples exposed to sidestream wastewater (DESeq2 ASVs; p_(adj)_ < 0.01), while another *Nitrosomonas* ASV dominated samples exposed to sidestream wastewater (Figure S3B).

**Figure 4:**
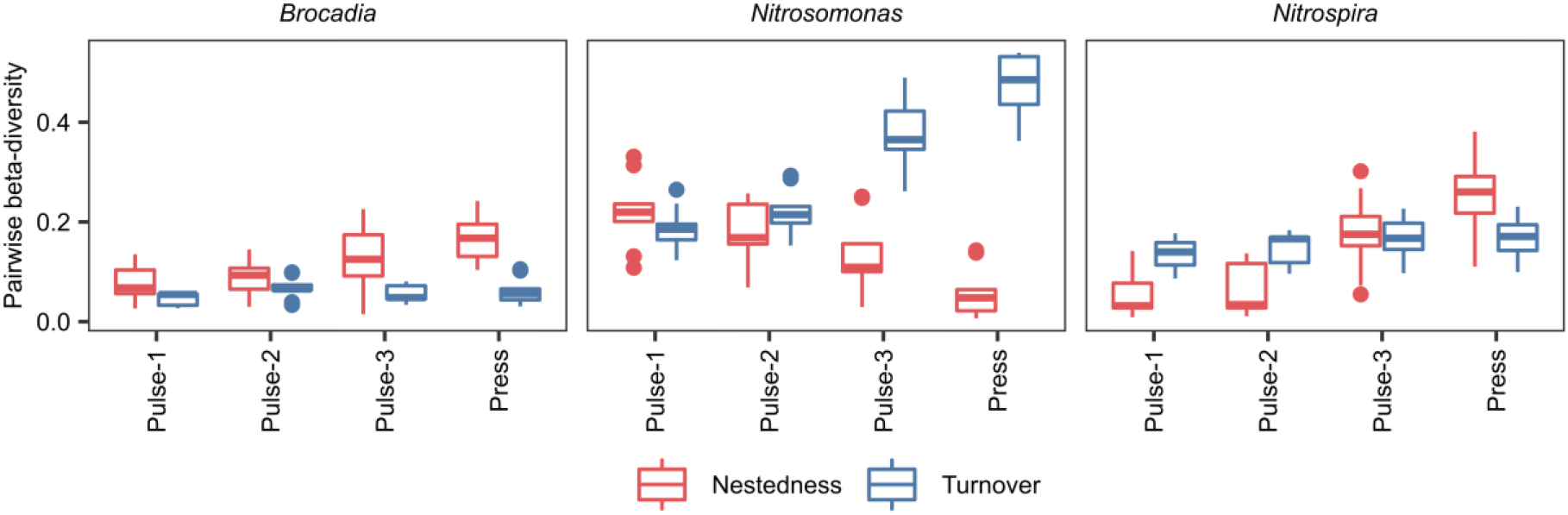
Pairwise beta-diversity between No-disturbance and other treatments using the Kaiju data from figure 3C. The presence-absence Sørensen index (q=0) was estimated as its two components: species turnover (i.e. the Simpson index) and dissimilarity due to nestedness.

To characterize the populations, metagenomic reads were assembled and resulting contigs were binned, resulting in 430 MAGs. Of these, five MAGs were affiliated to *Nitrosomonas* (Figure 5). Using a set of 203 single-copy genes, the taxonomic affiliation of four of these MAGs was established (Figure S4). The MAGs SJ624 and SJ754 belong to *Nitrosomonas* cluster 7, i.e. the *Nitrosomonas europaea / mobilis* cluster. While the MAG SJ624 is most closely related to *N. europaea*, SJ754 is most similar to *N. mobilis*, although it only shares 82.2% average nucleotide identity (ANI) (Figure S4). The MAG SJ76 is a member of *Nitrosomonas* cluster 6a (*Nitrosomonas oligotropha* cluster) sharing 99.7% ANI with *Nitrosomonas* sp. JL21, an AOB isolated from laboratory activated sludge ^60^. The MAG SJ634 is a member of *Nitrosomonas* cluster 6b, a group of AOB commonly associated with estuarine and marine environments ^61^. A fifth MAG, SJ328, was only 58% complete (File S1) and therefore was not included in this analysis. However, phylogenetic analysis of the ribosomal S2 gene indicates that SJ328 is closely related to SJ624 in cluster 7 (Figure S5). All five *Nitrosomonas* MAGs contained a mix of the essential genes required for both nitrification and for denitrification to nitrous oxide (N_2_O) (File S1).

**Figure 5.**
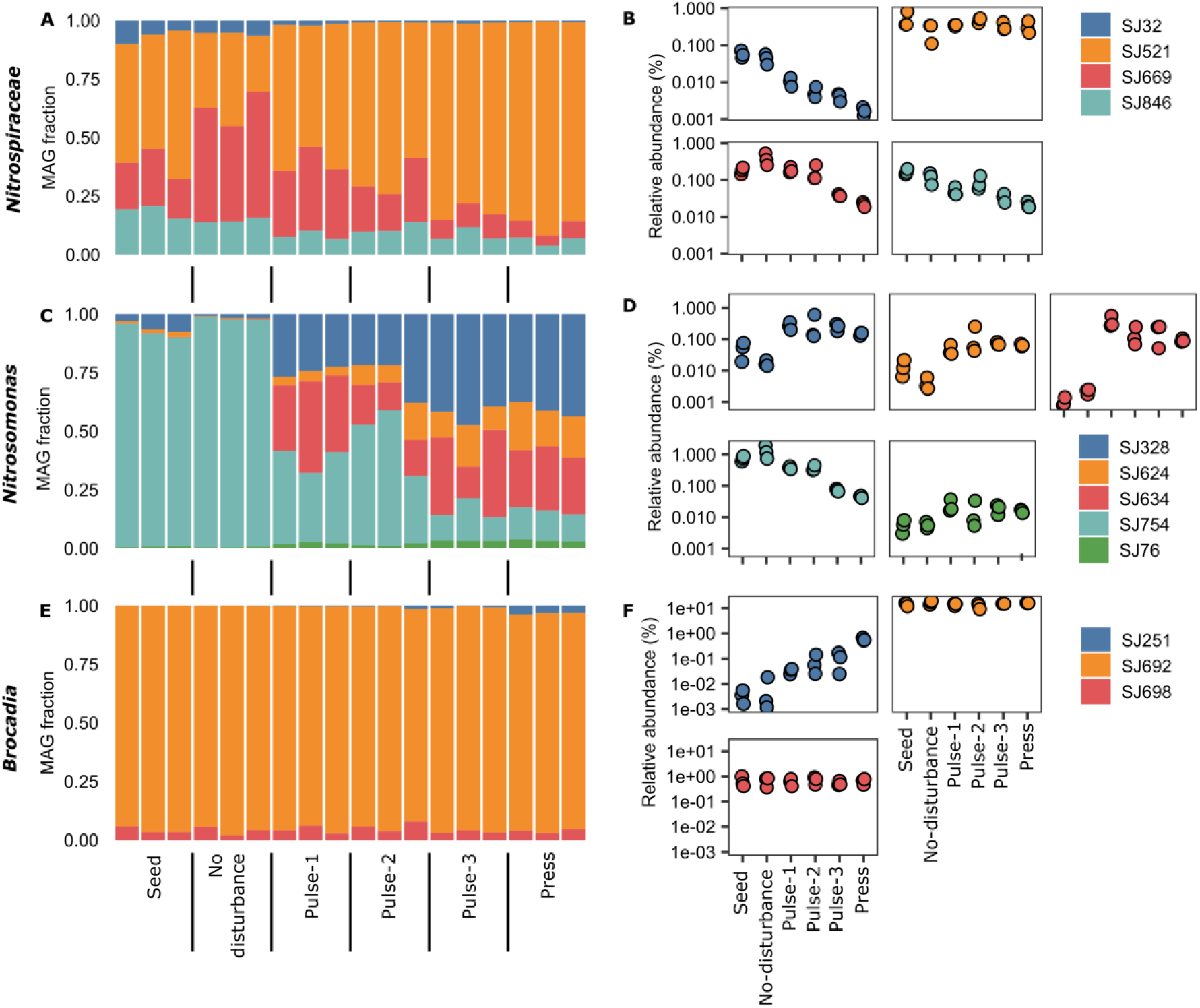
Abundance of MAGs classified as *Nitrospiraceae, Nitrosomonas* and *Brocadia*. **A, C, E:** Relative abundance within the group. **B, D, F:** Relative abundance against all the bins in the metagenome.

The MAG SJ754 made up more than 90% of the *Nitrosomonas* community in the mainstream (No disturbance) but decreased sharply in abundance with increasing sidestream exposure to about ∼13% at press disturbance (Figure 5C, D). Previously characterized *N. mobilis* sp. have been observed to be halotolerant and, in some cases, even obligately halophilic ^62,63^. Indeed, SJ754 contains several gene clusters that would facilitate an adaptation to saline environments (Supporting results and discussion, File S1). In addition, SJ754 is also the only AOB MAG to encode parts of both low and high affinity forms of RuBisCO, which may provide an advantage during times low and fluctuating CO2 concentrations. The concentration of inorganic carbon in the mainstream was more than tenfold lower than in the sidestream (Gustavsson et al. 2020), which may explain why SJ754 had a competitive advantage in the mainstream environment of fluctuating CO_2_ concentrations.

The other four *Nitrosomonas* MAGs were all present at higher relative abundances with increasing sidestream exposure. Interestingly, these AOB are expected to have very different optimal growth conditions based on their phylogeny ^61,64^. Being a member of the *N. oligotropha* cluster 6a, SJ76 represents an AOB most likely adapted to low ammonium concentrations, in contrast to the other four *Nitrosomonas* MAGs. Hence, alternative metabolisms may be an important factor to explain the continuous low relative abundance of SJ76. Indeed, SJ76 and SJ634 both contain genes necessary for alternative metabolisms such as urea and hydrogen utilization. The closely related SJ624 and SJ328, belonging to *Nitrosomonas* cluster 7, both encode a possible high affinity terminal oxidase (sNOR), shown to be upregulated in AOB grown under oxygen-limiting conditions ^65^. Lastly, SJ634, which belongs to *Nitrosomonas* cluster 6b, encodes a variety of genes related to halotolerance, including: 1) a sodium motive force driven type III NAD dehydrogenase, 2) a sodium motive force driven N-type ATPase ^66^, and 3) biosynthesis genes for the compatible solutes ectoine and hydroxyectoine ^67^ (File S1). These differences could lead to niche differentiation between the AOB and perhaps explain the observed AOB turnover.

Alternatively, or concurrently, fluctuations in abundance of AOB MAGs could be due stochastic processes rather than niche differentiation ^19^, as migration of AOB from upstream to downstream processes is known to occur in WWTPs ^68^. Thus, the high abundance of SJ754 in the mainstream samples could have been caused by continuous immigration from the upstream activated sludge, akin to the mass-effect perspective in the metacommunity framework ^69^. However, given the low sludge retention time (2-3 days) in the preceding high-rate activated sludge with no signs of nitrification in the investigated period, such stochastic effects were likely unimportant for both the *Nitrosomonas* and the *Nitrospira* communities

### A lineage I *Nitrospira* was resistant to sidestream exposure

Four *Nitrospira* MAGs were resolved after binning (Figure 5A). Three of them are members of *Nitrospira* lineage I (SJ521, SJ669, and SJ846). The fourth is a member of *Nitrospira* lineage II (SJ32), a group that comprises both nitrite oxidizers and comammox bacteria (Figure S7) ^29,32^. Although comammox *Nitrospira* have been observed in WWTPs ^70,71^, a search for *amoA* in the entire metagenome assembly did not identify any *Nitrospira amoA*, suggesting that comammox *Nitrospira* were not present or were below detection. For more details, see Supplementary results and discussion.

In general, the *Nitrospira* populations were sensitive to sidestream exposure, with the relative abundance of three of the four detected MAGs decreasing in abundance with increasing sidestream exposure (Figure 5B). These results align with the observed decrease in *Nitrospira* alpha-diversity (Figure 3). The sole exception was SJ521, a *Nitrospira* lineage I NOB, whose relative abundance remained the same regardless of sidestream exposure (Figure 5B).

As SJ521 was the dominant *Nitrospira* MAG in Press-disturbance, it was hypothesized that this MAG would encode genes providing this species with niche differentiating traits absent in the other *Nitrospira* and thus a competitive advantage. Previously, several *Nitrospira* species have been shown to possess genes that enable use of alternative energy sources such as ammonium, formate, or hydrogen and are able to engage in reciprocal feeding with ammonia oxidizers through the turnover of urea or cyanate ^26-28^.

Three of the four *Nitrospira* MAGs, including SJ521, encode a very similar suite of genes for these alternative metabolisms (File S1). However, the dominant MAG SJ521, is the only one that contains a Na^+^/H^+^ antiporter complex, which can regulate cytoplasmic pH and/or Na^+^ concentration and confer halotolerance ^72^. To date, the only *Nitrospira* genomes observed to encode this kind of antiporter complex have been found in highly saline / alkaline environments ^73^. Interestingly, similar Na^+^/H^+^ antiporter complexes were also found in four of the five *Nitrosomonas* MAGs (File S1).

The presence of halotolerant populations of *Nitrospira* and *Nitrosomonas* after sidestream exposure is unexpected, as the concentrations of Na^+^ ions were the same in the mainstream and sidestream (Table S1). Nonetheless, the conductivity, which is a measure of the total concentration of ions in the water, was more than five-fold higher in the sidestream. This is an indication of higher osmotic stress in the sidestream. In this environment, adaptations to osmotic stress might offer a competitive advantage to halotolerant populations. Hence, it is possible that uncharacterized adaptations to osmotic stress in *Nitrospira* SJ521 could explain the persistence of this population.

### Abundance of *Brocadia fulgida* increased with sidestream exposure

Three *Brocadia* MAGs were present in the PNA biofilms (Figure 5E). SJ692 was the dominant anammox MAG in the dataset, and its abundance did not change with sidestream exposure. It had 98.9% ANI to *Brocadia sapporoensis*, which had previously been identified as the dominant anammox bacterium at the Sjölunda PNA bioreactors ^44^, as well as in other wastewater bioreactors ^74,75^. Another *Brocadia* MAG with the same abundance across samples was SJ698, which appears to be a new species of *Brocadia* (Figure S8), although its 16S gene had a 98% identity to a sequence from the Yangtze estuary in China ^76^. For more details, see Supplementary results and discussion.

SJ251 had its highest abundance in the Press-disturbance samples (Figure 5F), which explains the increase in functional diversity of anammox genes with sidestream exposure (Figure 3C). The highest ANI for SJ251 was 99.2% to the *B. fulgida* genome. All the anammox MAGs had an acetyl-CoA synthetase, which would allow converting acetate into acetyl-CoA. However only SJ251 encoded genes for acetate kinase (*ackA*) and phosphate acetyltransferase (PTA), where acetyl-CoA is made via acetyl phosphate. In *Escherichia coli, ackA* + PTA are used at high acetate concentrations ^77^, and this agrees with a previous report of *B. fulgida* being enriched when acetate was added to the medium ^78^. The sidestream reactor was located downstream of anaerobic biodigesters with considerable effluent concentrations of volatile fatty acids (VFA) (196 mg COD/L). This was much higher than the concentrations of VFA in the feed to the mainstream reactors (< 20 mg COD/L). Due to the availability of acetate and other VFA, SJ251 might have a competitive advantage that differentiates their niche from the other anammox bacteria and allows them to prosper in the sidestream.

### Genes for denitrification were ubiquitous in the PNA biofilms

As commonly observed in microbial communities ^37,79,80^, few genomes in the PNA biofilms carried genes for the full denitrification pathway, although genes for partial denitrification were common (Supporting results and discussion, Figure S9, S10).

To test if sidestream exposure had an effect on the abundance of denitrifiers, we estimated the overall abundance of MAGs having the genes *nirK, nirS, norB* and *nosZ*. The total relative abundance was the same across the disturbance regimes (Figure S11), although shifts in abundance occurred within the distribution of MAGs (Figure S12-S15). This variation at the individual sample level but stability at a broader level signifies functional redundancy and is described by the portfolio effect ^81^. Since denitrification is a trait with a wide phylogenetic distribution, is likely to be more resistant to disturbances than traits with a narrow phylogenetic distribution like nitrification and anammox.

### Links in abundance patterns suggest interactions among taxa in the biofilms

Since changes in relative abundance were observed for ASVs and MAGs across samples (Figure 2), we used SPIEC-EASI^82^ to find potential links in abundance between both ASVs (Figure S3) and MAGs (Figure 6). In both the ASVs and the MAGs networks, *Nitrosomonas* ASVs and MAGs were part of distinct clusters, representing populations either sensitive or resistant to sidestream exposure. It is also noteworthy that the abundance of *Nitrosomonas* SJ754, was linked to *Nitrospira* SJ669 and SJ846 (Figure 6). Both might be equally affected by sidestream exposure; the link could also indicate syntrophy, with *Nitrospira* using the NO_2_^-^ produced by *Nitrosomonas*, or reciprocal feeding by *Nitrospira* converting urea or cyanate to ammonia. If they are co-dependent, inhibition of *Nitrosomonas* SJ754 alone, might explain the decrease in abundance of *Nitrospira* SJ669 and SJ846 or vice versa. This may have importance for successful management of PNA biofilms.

**Figure 6.**
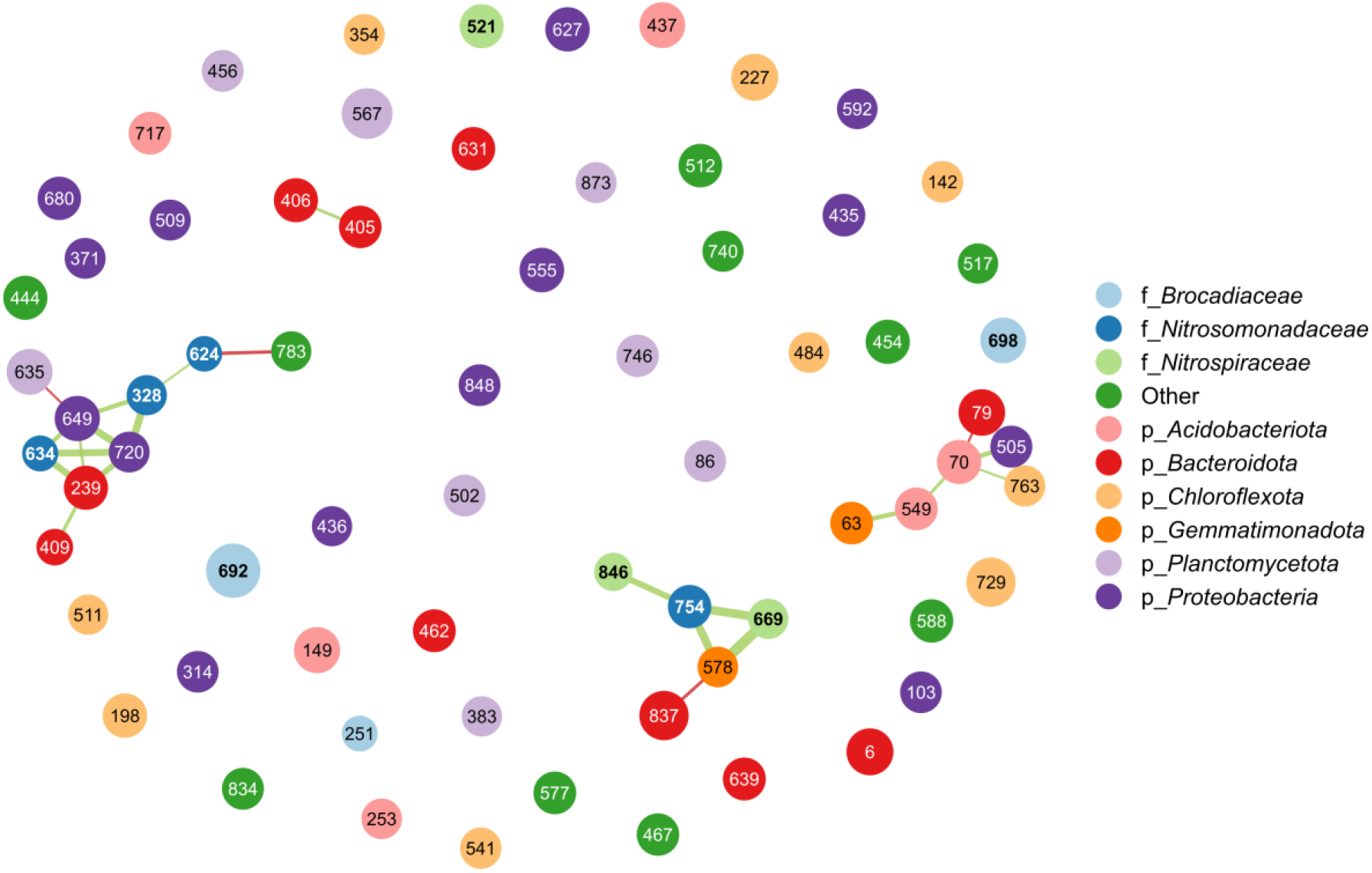
Network analysis of abundant MAGs. Each circle (node) represents a MAG; connections (edges) indicate that MAG abundances are not independent of each other. Green edges show a positive link, while red edges indicate a negative link. The size of the node is proportional to its mean centred-long-ratio abundance. The thickness of an edge indicates edge weight.

The abundance of other *Nitrosomona*s ASVs and MAGs (SJ328, 624, 634) was linked to the abundance of ASVs and MAGs within *Bacteroidota* and *Proteobacteria* in a different cluster (Figure 5, Figure S3). It is tempting to suggest that the latter taxa were denitrifying, as indicated by the presence of genes for denitrification in the Proteobacterial MAGs (Figure S9, S10), using NO_2_^-^ produced by the *Nitrosomonas*. Alternatively, carbon fixed by the *Nitrosomonas* may have been used for heterotrophic growth ^83-85^.

## CONCLUSIONS

This study shows that it is possible to manipulate the microbial community of a mature biofilm towards a desired state. With increased sidestream wastewater exposure, the functional versatility of PNA biofilms, and hence their functional redundancy, decreased for the unwanted functional group (NOB), but increased for the wanted functional groups (AOB and anammox bacteria). The observed responses to the disturbances demonstrate that exposure of sidestream wastewater has potential as a tool to regulate PNA biofilm communities. Our metagenomic analyses show that the pangenomes of nitrifiers and anammox bacteria included genes for osmotic stress response, organic matter and inorganic nitrogen usage. This provides not only multifunctionality but also functional redundancy, and thus resistance to the loss of services and disservices in ecosystems. For WWTPs, our findings imply that knowledge about the versatility of pangenomes should be considered when designing long-term operational strategies. Although we observed a range of effects of sidestream exposure on the *Nitrosomonas, Nitrospira*, and *Brocadia* populations, the detailed mechanisms remain unclear. Future studies using other approaches, like metatranscriptomics and activity measurements, may shed further light on the stress response to sidestream exposure.

## METHODS

### Study design

Two parallel PNA MBBRs at the Sjölunda WWTP (Malmö, Sweden) were fed with either mainstream wastewater from a high-rate activated sludge plant for organic carbon removal or sidestream water, sludge liquor from anaerobic sludge digestion. Both bioreactors were filled with biofilm carriers (K1®, Veolia Water Technologies AB – AnoxKaldnes, Lund, Sweden), which offer a protected biofilm surface. The reactors are described in detail elsewhere ^44^. Conditions in the bioreactors are shown in Table S3.

The biofilm carriers were kept isolated from other carriers in the MBBRs in cylindrical cages. A steel mesh bottom of the cages allowed water circulation (immersed volume 2.5 L, 30% filling). The cages were filled with biofilm carriers on November 11, 2015 and sampled on January 7, 2016. Samples were also taken from the mainstream PNA MBBR at the start of the experiment on November 11. These samples are referred as Seed.

### Sequencing and data analysis

Amplicon sequencing of the 16S rRNA V4 region was conducted using a MiSeq (Illumina), with 12 biofilm carriers from each treatment. For each treatment, three samples were selected for metagenomics based on shotgun sequencing using the NovaSeq 6000 platform (Illumina). Details of sampling, amplicon sequencing, shotgun metagenomics, and data analysis are described in the supporting information.

## DATA AVAILABILITY

Amplicon sequencing reads, raw shotgun metagenomics reads, and metagenome assembled genomes (MAGs) are available at NCBI under the bioproject PRJNA611787. All data generated or analysed during this study will be available upon request to the corresponding author.

## AUTHOR CONTRIBUTIONS

C.S., F.P. and M.H. designed the experiment. C.S. did the analysis of metagenome dataset. C.J.S. analysed the *Nitrosomonas* and *Nitrospira* genomes. D.G. operated the reactors. O.M. performed the salinity measurements. A.E. contributed to the interpretation of the results. All authors contributed to the writing of the manuscript. All authors reviewed and approved the final manuscript.

## ACKNOWLEDGEMENTS

This work was funded by FORMAS (contract no. 2018-01423). The authors acknowledge the support from the National Genomics Infrastructure in Stockholm funded by Science for Life Laboratory, the Knut and Alice Wallenberg Foundation and the Swedish Research Council, and SNIC/Uppsala Multidisciplinary Center for Advanced Computational Science for assistance with massively parallel sequencing and access to the UPPMAX computational infrastructure. Assembly and binning were performed on resources provided by SNIC through UPPMAX under the projects SNIC 2020-15-14 and 2020-16-18. We also acknowledge the colleagues at the Sjölunda WWTP, for monitoring the pilot reactors.

## COMPETING INTERESTS

The authors declare no competing interests.

## Notes

### Competing Interest Statement

The authors have declared no competing interest.

