## Supporting information for "Disturbance-based management of ecosystem services and disservices in partial nitritation anammox biofilms"

#### SUPPORTING METHODS

##### Biofilm sampling

For each treatment, twelve biofilm carriers were snap-frozen in an ethanol-dry ice mixture immediately at sampling, kept frozen in dry ice during transportation and then stored at -80°C. The biofilm was removed from the carrier compartments and added to a lysis matrix tube E (MP biomedicals, Santa Ana, CA, USA) with 800 µl of lysis solution of a ZR-duet MiniPrep kit (Zymo Research). Mechanical disruption of the biofilm was done with a FastPrep-24 5G (MP biomedicals) at speed 6 for 40 seconds. Subsequent steps of the DNA extraction were done with the ZR-duet kit according to manufacturer instructions.

##### 16S sequencing

PCR amplification of the 16S rRNA V4 region (16S) was done with primers 515′F [1] and 806R [2], using dual indexing of the primers [3]. PCR products were purified with Ampure XP (Beckman Coulter, Brea, CA, USA), and PCR amplicons were then pooled in equimolar amounts. Sequencing was done on an Illumina MiSeq using the MiSeq Reagent Kit v3 (Illumina, San Diego, CA, USA). Three samples from Pulse-1 with less than 30.000 reads were excluded prior to analysis. DADA2 version 1.16 [4] was used to infer amplicon sequence variants (ASVs), with settings *pool=TRUE*. SILVA 138 [5] was used for taxonomic classification of the 16S amplicons.

##### Shotgun metagenomics

For each treatment, three samples were selected for shotgun metagenomics. Libraries were prepared with a TruSeq PCR-free kit (Illumina). Samples were then sequenced on a NovaSeq 6000 with a 2x151 setup, resulting in 800 million pair-reads. Low quality reads and adapters were removed with Trimmomatic v0.39 [6]. Co-assembly of short reads was done with Megahit [7] using the --*presets meta-large* setting, resulting in 1,975,712 contigs of $\geq$1 kb, and N50 of 4375 bp. Reads were then mapped to the co-assembly using Bowtie2 v2.3.5 [8]. Protein coding sequences in all the assembled contigs were identified with Prodigal v.2.6.3 [9], using the meta prediction mode.

**Diversity analysis**

SingleM (https://github.com/wwood/singlem) was used to create an OTU-like table of 14 single copy marker genes, which was used to estimate alpha and beta-diversity in the metagenomics dataset. After subsampling to even depth alpha diversity was estimated as species richness for both amplicon sequencing variants (ASVs) based on 16S amplicon sequencing and metagenomics marker genes. Because PERMANOVA is sensitive to unbalanced designs, three samples were randomly excluded from each group in the 16S dataset, except for Pulse-1. Thus each group in the 16S dataset had nine samples.

Using the R package hilldiv [10], dissimilarity indices were calculated based on partitioning of Hill numbers [11] with a Sørensen-type overlap for both the 16S dataset and the metagenomics one. Beta-diversity was estimated from q=0 to q=3 in 0.1 intervals. Among samples taken at day 58, Permutational multivariate analysis of variance (PERMANOVA) [12] was used to test for significant difference between group centroids.

Kaiju [13] was used for taxonomic classification of assembled genes against the NCBI BLAST nr database. Richness (number of classified genes) was estimated for *Nitrosomonas*, *Nitrospira* and *Brocadia*. Using the kaiju data, the presence-absence Sørensen index, and its two components, turnover and dissimilarity due to nestedness [14] were also estimated. To identify gene clusters, the proteins classified as *Nitrosomonas*, *Nitrospira* and *Brocadia* were aligned against each other with DIAMOND [15] using --ultra-sensitive parameter, followed by clustering with MCL [15], with an inflation factor of 8.

Hidden Markov models (HMM) from the FunGene database [16] were used to search for AOB, *Nitrospira* or Archaea *amoA* in the metagenomics assembly with HMMER 3.3 (hmmer.org). The lowest E-value and highest bit score always corresponded to the amoA_AOB profile. In addition, the recovered *amoA* sequences were analysed with an LG+F+G4 phylogenetic tree, showing that all *amoA* corresponded to known *Nitrosomonas amoA* clusters (Figure S6). PCR with comammox *amoA* primers [17] led to nonspecific PCR amplification, as determined by sequencing of the PCR product.

**Metagenome assembled genomes**

Metagenomic binning was done in MetaBAT2 [18], resulting in 877 bins. Completeness, contamination, and relative abundance of the bins was estimated with CheckM [19]. 430 metagenome assembled genomes (MAGs) had more than 50% completion and less than 10% contamination. Taxonomic classification of these MAGs was done using GTDB-Tk [20] with the GTDB r89 taxonomy [21,22]. To study evolutionary relationships of the *Nitrosomonas*, *Nitrospira* and *Brocadia* MAGs, the MAGs were compared with genomes from the NCBI assembly database. Phylogenomic trees based on single-copy genes were made with GtoTree v1.2.1 [23], using the Betaproteoabacteria HMM-set (203 genes) for *Nitrosomonas*, and the Bacteria HMM-set (74 genes) for *Nitrospira* and *Brocadia*. The multiple sequence alignment from GtoTree was used to generate maximum likelihood phylogenetic trees in IQ-TREE v.2.0.3 [24] with 1000 rapid bootstrap replicates; a substitution model for each gene in the alignment was chosen with ModelFinder [25] using the partition files generated by GtoTree. Average nucleotide identity (ANI) estimations were done with FastANI [26]. The online version of eggNOG-mapper [27,28] was used to annotate the *Nitrosomonas*, *Nitrospira* and *Brocadia* genomes. The *Nitrosomonas* and *Nitrospira* genomes were compared to each other and against previously sequenced nitrifiers using the integrated microbial genomes (IMG) database and the comparative analysis system [29], as well as Genoscope [30]. With HMMER we searched for the presence of the genes *amoA*, *narG*, *napA*, *nirA*, *nirB*, *nrfA*, *nirS*, *nirK*, *norB* and *nosZ* in all the MAGs. Using the bacterial phylogenetic tree generated by GTDB-Tk, phylogenetic signal of those traits was estimated as the D statistic [31] with the R package caper [32].

**Network analysis**

Using the relative abundance of the 16S ASVs or MAGs, network analysis was carried out using the R package SPIEC-EASI [33] with the glasso method (lambda.min.ratio=.01, nlambda=30, rep.num=100). Clusters in the network were identified with the walktrap community finding algorithm.

**Salinity measurements**

Three water samples from sidestream and mainstream at the Sjölunda WWTP were collected in January 2021. Conductivity was measured using a CO11 conductivity sensor (VWR, Radnor, Pennsylvania, United States). The concentration of $Na^+$, $Cl^-$, $NH_4^+$, $K^+$, $PO_4^{3-}$, $Mg_2^+$, $Ca_2^+$ and $SO_4^{2-}$ was measured with ICS-900 ion chromatographs (Dionex Sunnyvale, California, USA).

### SUPPORTING RESULTS AND DISCUSSION

***Nitrosomonas* MAGs**

The AOB SJ754 encodes two forms of complex I (NADH:quinone oxidoreductase or NADH dehydrogenase) of the electron transport chain. It encodes a canonical type I NADH dehydrogenase (*nuoA-N*), parts of which are found in all five AOB MAGs and a sodium dependent NADH:quinone oxidoreductase (type III NADH dehydrogenase), which can regulate the cytoplasmic sodium concentration. If operated in reverse, this complex would allow for the generation of NADH with sodium motive force [34], providing an energetic advantage over AOB competitors that depend solely on proton motive force to generate cellular reducing power.

Although none of the *Nitrosomonas* MAGs included an *amoA* gene, multiple *amoA* sequences were found in the metagenomic assembly, corresponding to *Nitrosomonas* cluster 7, 6a, 6b and 8 (Figure S6). The absence of *amoA* in the MAGs, might be due to the presence of multiple copies of *amoA* being present in the genome, as seen for *N. europaea*, [35], which makes difficult binning.

***Nitrospira* MAGs**

All four *Nitrospira* MAGs contained the expected genes essential for nitrite oxidation, and genes indicating a reverse TCA cycle method of carbon fixation (File S1) [36]. Although MAG SJ32 encodes many of the genes necessary for a possible functional hydrogenase, the relative abundance of SJ32 in the *Nitrospira* population was low in all treatments and decreases with increased sidestream exposure (Figure 5). Therefore, it seems that this does not provide a competitive advantage in these environments. In addition, SJ669 does not encode for all genes encoding any of these alternative metabolisms, but this is most likely due to the incompleteness of the MAG (~58%) (File S1).

**Anammox MAGs**

*B. sapporoensis* is a comparably fast growing anammox bacterium [37]. It has been proposed that the small genome size of *B. sapporoensis* is linked to its fast growth [38]. A genome size of 2.9 Mb was reported for *B. sapporoensis* [38], while a 2.8 Mb genome was observed for SJ692, with a 95% completion as estimated with CheckM (File S1). This was in fact smaller than two other two anammox MAGs in the PNA biofilm which both had a genome size of 3.2 MB and were estimated to be >95% complete with CheckM (File S1).

One mechanism that could allow SJ698 to compete with the dominant SJ692 is the presence of a Type VI secretion system (T6SS) in the SJ692 genome. T6SSs are used by bacteria to export proteins to neighbouring cells [39]. Thus the T6SSs in the anammox SJ698 could be used against other bacteria [40] or might provide protection against phagocytic amoebae [41]. Selective protozoan predation of anammox bacteria has been reported before [42], and hence it is possible that anammox bacteria differ in their defence strategies against predation. In addition, we observed a cyanase in the SJ698 genome. Cyanases are uncommon among anammox bacteria, with the exception of some marine anammox bacteria in the genus *Scalindua* [43,44]. This could provide an alternative source of ammonium to SJ698.

**Denitrifiers**

There appears to be coherence for some traits at high phylogenetic ranks, with the strongest signals occurring for *nirK*, *nirS* and *nosZ* (Table S2). For example, *nosZ* was observed in 59 out of 73 *Bacteroidota* MAGs, 15 of 17 *Myxococcota* (deltaproteobacteria) MAGs and 8 of 10 *Gemmatimonadota* MAGs, confirming reports of *Gemmatimonadota* being a potential sink for $N_2O$ [45,46]. Because of their abundance, *Bacteroidota* might represent the main $N_2O$ reducer in the PNA biofilms; the most abundant one being SJ837, a MAG classified as genus UTBCD1 (Figure S9, S10), which has been observed before in anammox granules [47] where it was also proposed to be an $N_2O$ reducer. Nitrification by AOB is often linked to $N_2O$ emissions [48]; bacteria with a *nosZ* gene might reduce $N_2O$. The potential to reduce NO was present in 20 out of 38 *Acidobacteria* MAGs, 48 of 105 *Proteobacteria* MAGs and 27 of 46 *Planctomycetota* MAGs. With a mean abundance of 3%, SJ567, a *Planctomycetota* classified as UTPLA1, might be the main NO reducer (Figure S9, S10), but as it lacks a *nosZ gene* it is likely an $N_2O$ producer.

### SUPPORTING FIGURES

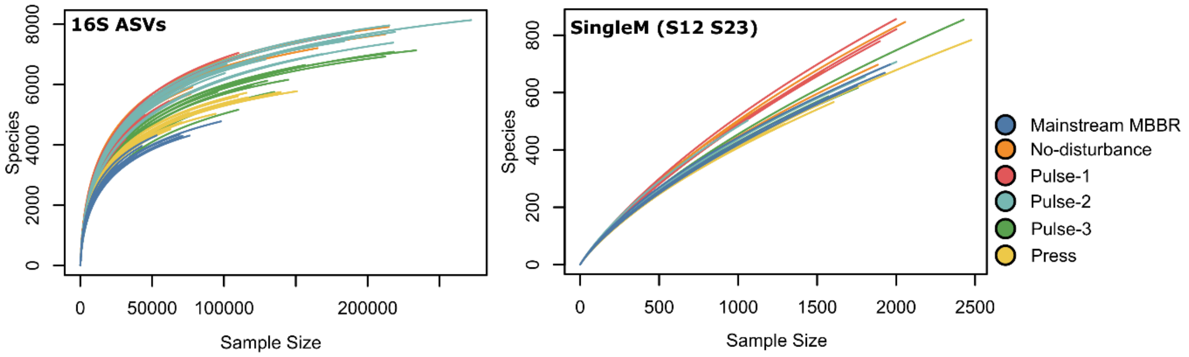

**Figure S1:** Rarefaction curve for 16S ASVs and the metagenomics dataset

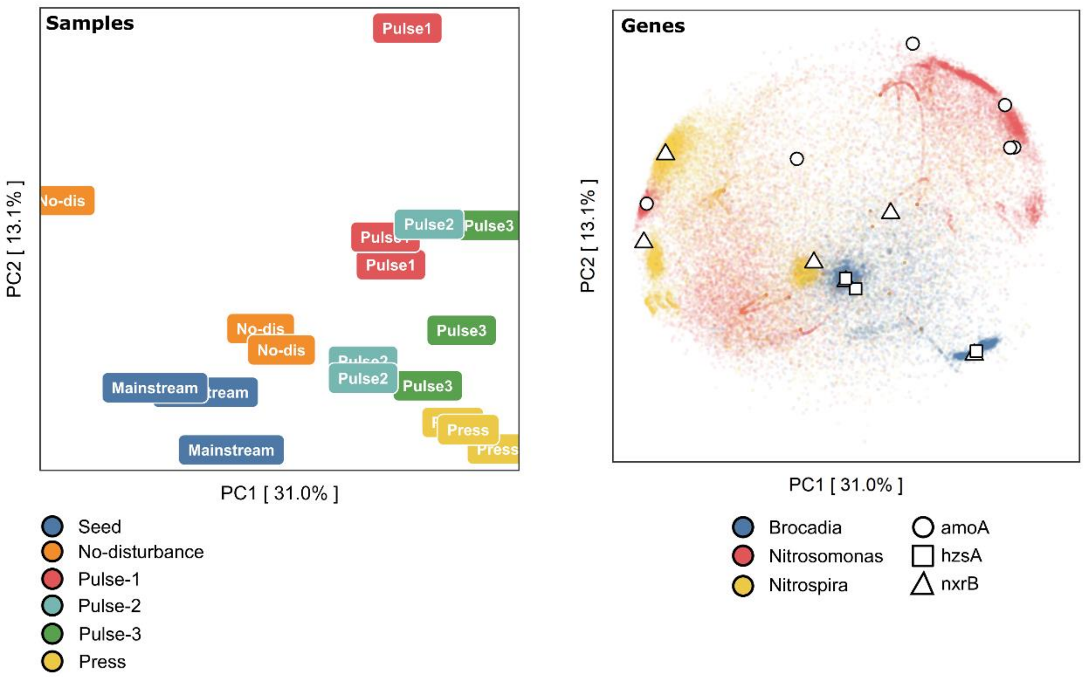

**Figure S2:** PCA biplot of *Nitrosomonas*, *Brocadia* and *Nitrospira* genes using the kaiju data; scores are shown in the left and loadings at the right. **Left:** The labels indicate biofilm samples (scores). **Right:** Each dot indicates an anammox or nitrifier gene (loadings); *amoA*, *hzsA* and *nxrB*/*narG* are shown as circles, squares, and triangles respectively.

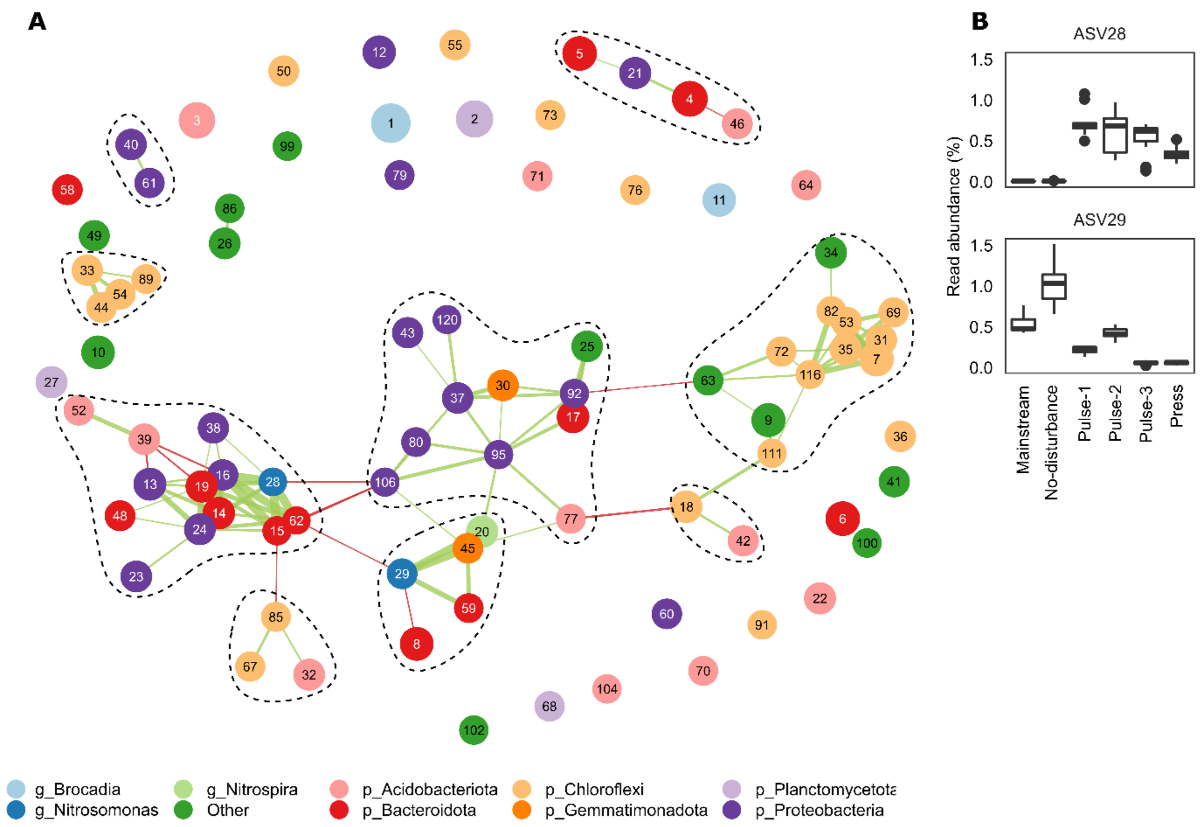

**Figure S3. A:** Network analysis of abundant 16S ASVs. Each circle (node) represents an ASV; connections (edges) indicate that ASV abundances are not independent of each other. Green edges show a positive link, while red edges indicate a negative link. The size of the node is proportional to its mean CLR abundance. The thickness of an edge indicates edge weight. **B:** Read abundance of the two most abundant *Nitrosomonas* ASVs.

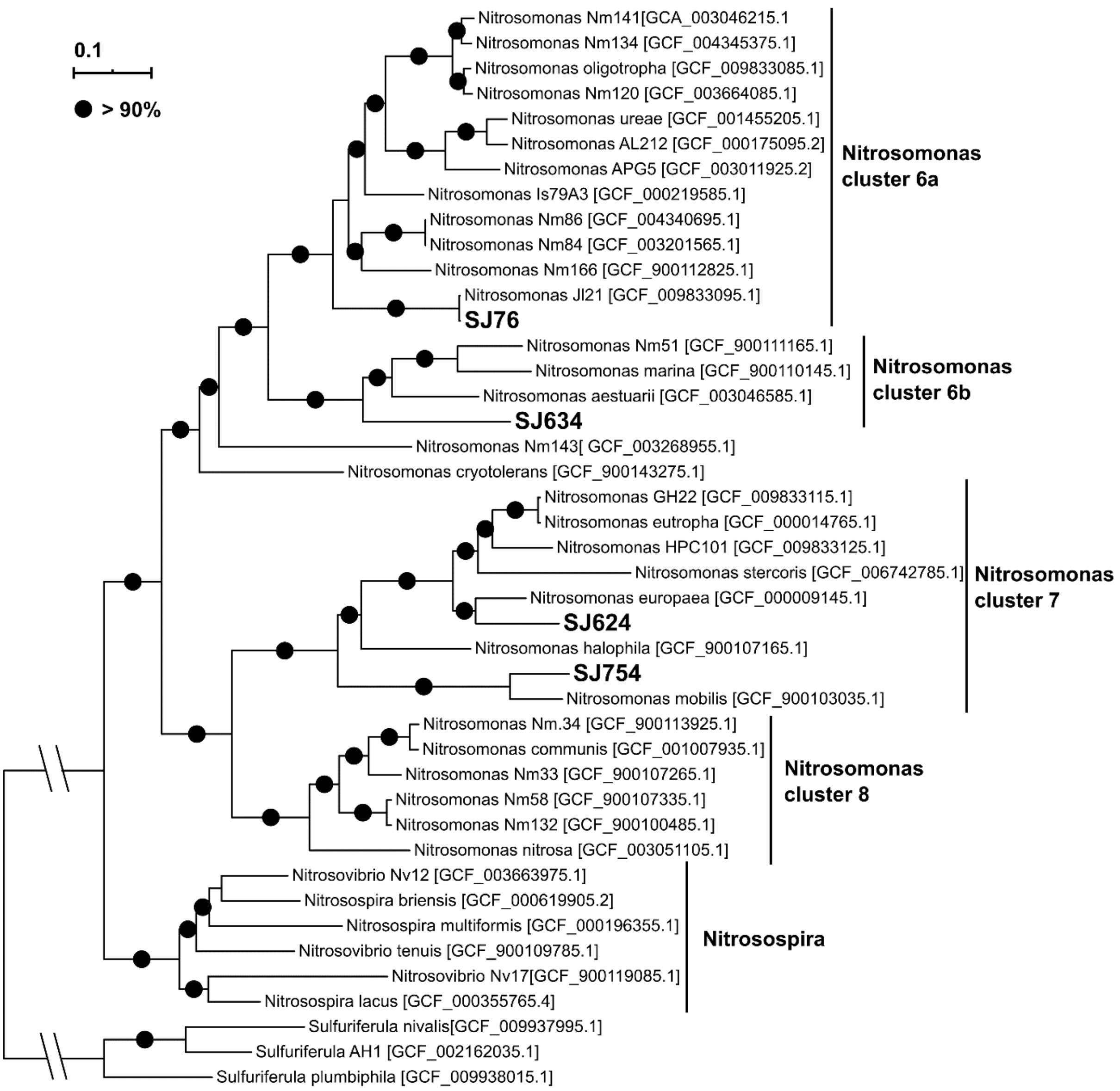

**Figure S4:** Phylogenetic tree for MAGs classified as *Nitrosomonas* based on 203 single-copy genes. Circles show > 90% ultrafast bootstrap support for a clade. *Sulfuriferula* was used as outgroup.

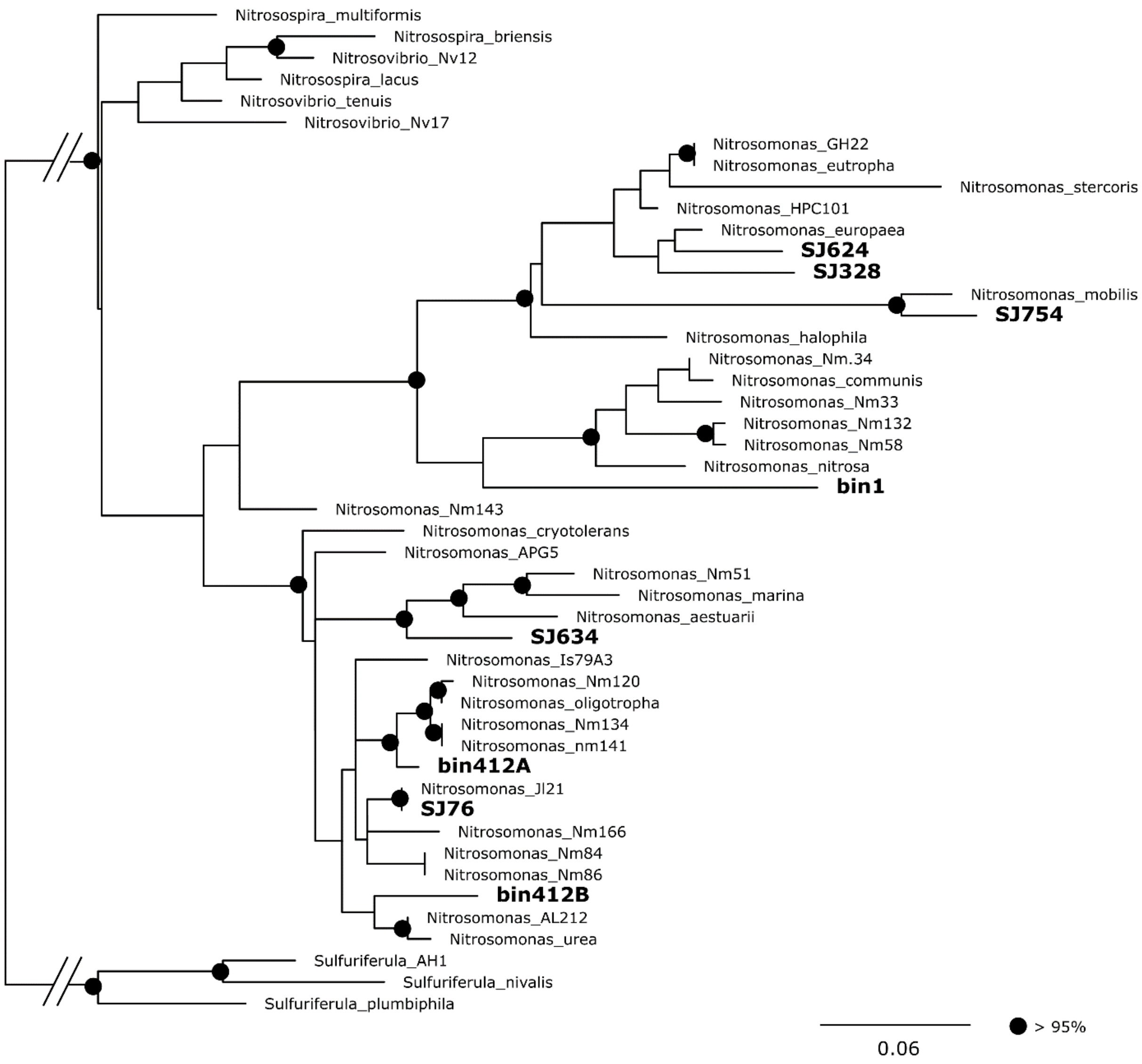

**Figure S5:** Phylogenetic tree of ribosomal protein S2 in MAGs classified as *Nitrosomonas*, using a LG+R3 substitution model. Circles show >= 95% ultrafast bootstrap support for a clade. *Sulfuriferula* was used as outgroup. Bin412 is a bin with high contamination, and bin1 is a bin with low completion.

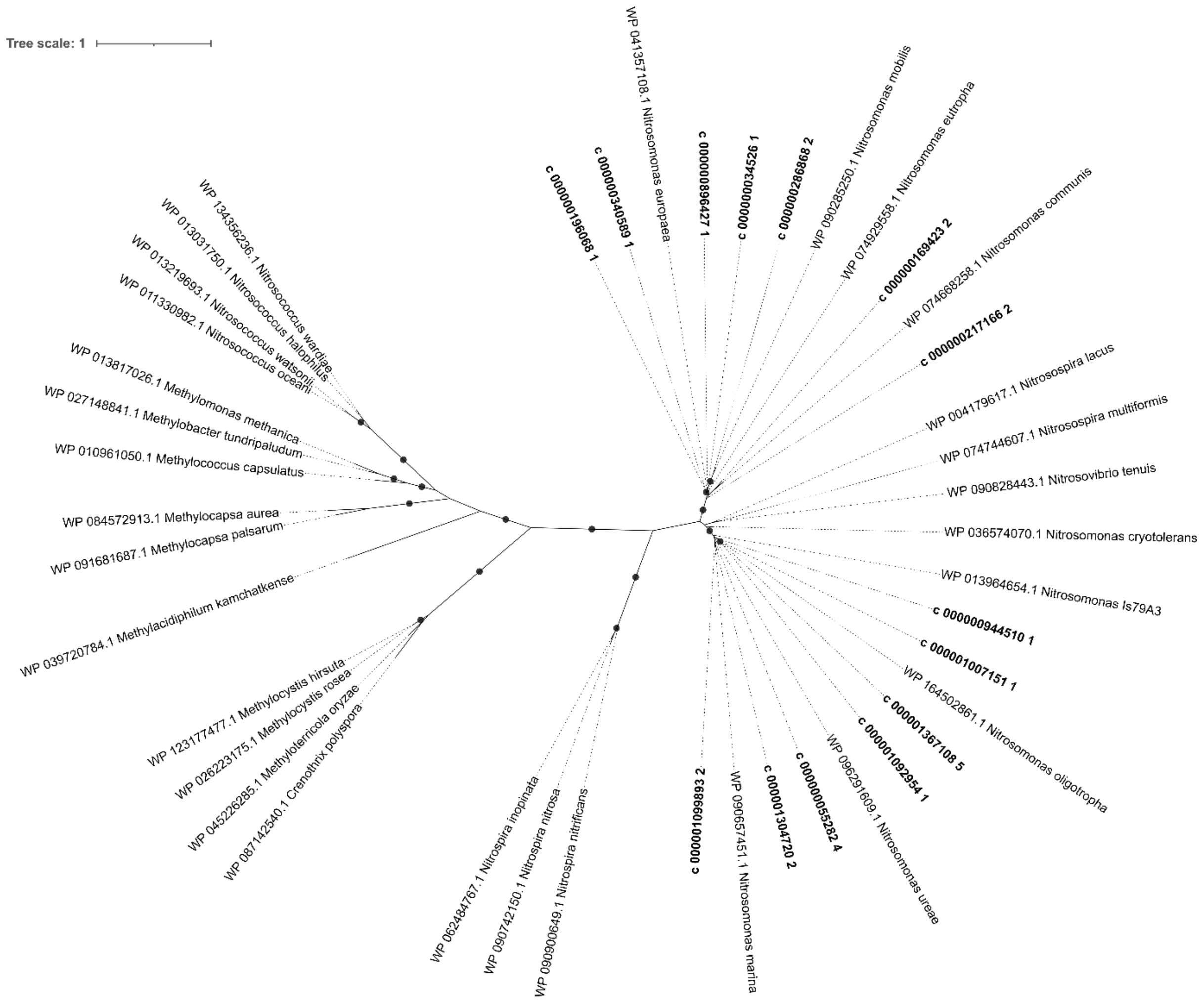

**Figure S6:** Unrooted phylogenetic tree of the bacterial *amoA* and *pmoA* gene. Labels in bold indicate *amoA* sequences from this study. Circles show >= 95% ultrafast bootstrap support for a clade.

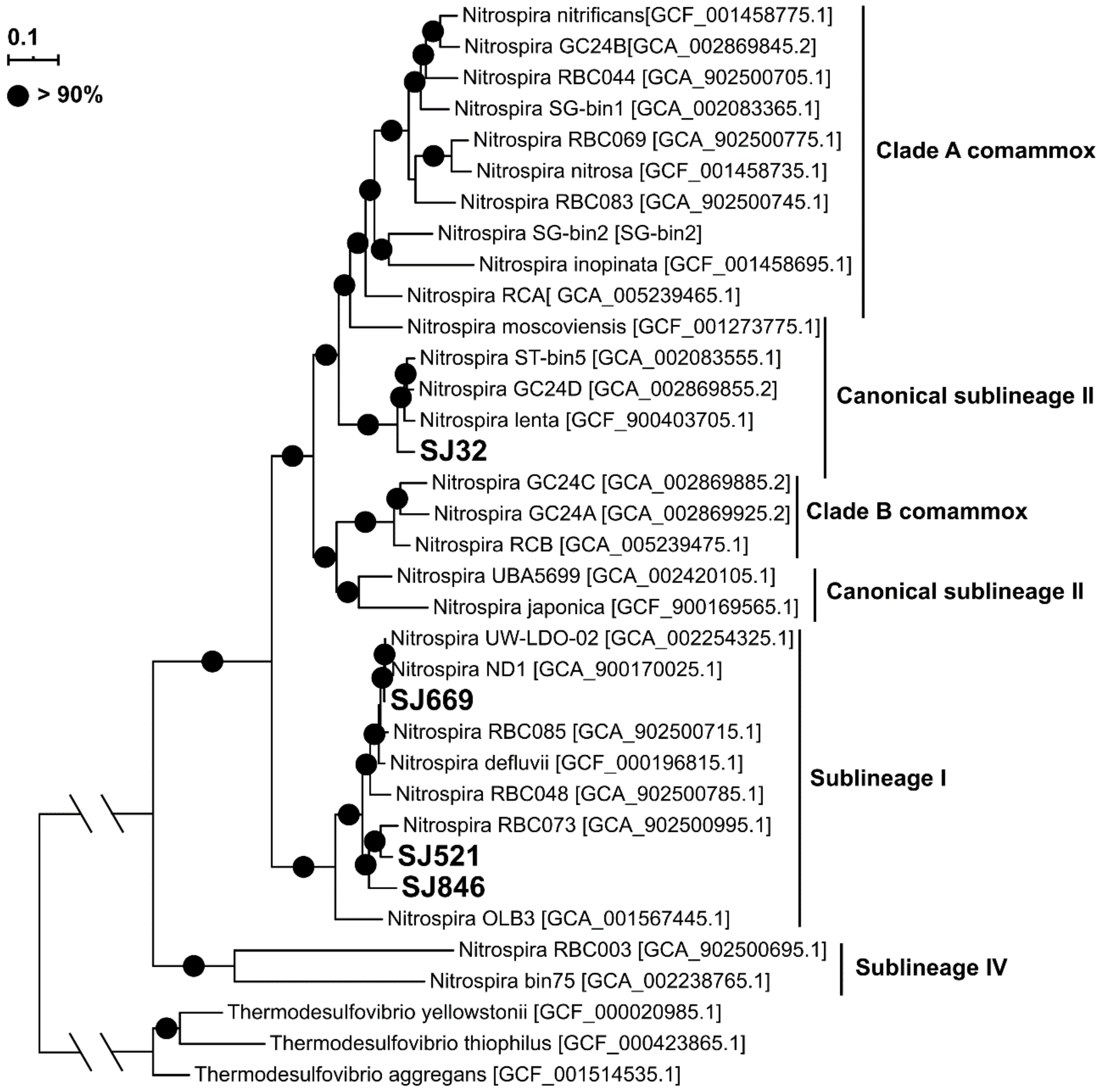

**Figure S7:** Phylogenetic tree for MAGs classified as *Nitrospira* based on 79 single-copy genes. Comammox clades were identified based on literature. *Thermodesulfovibrio* was used as outgroup. Circles show > 90% ultrafast bootstrap support for a clade.

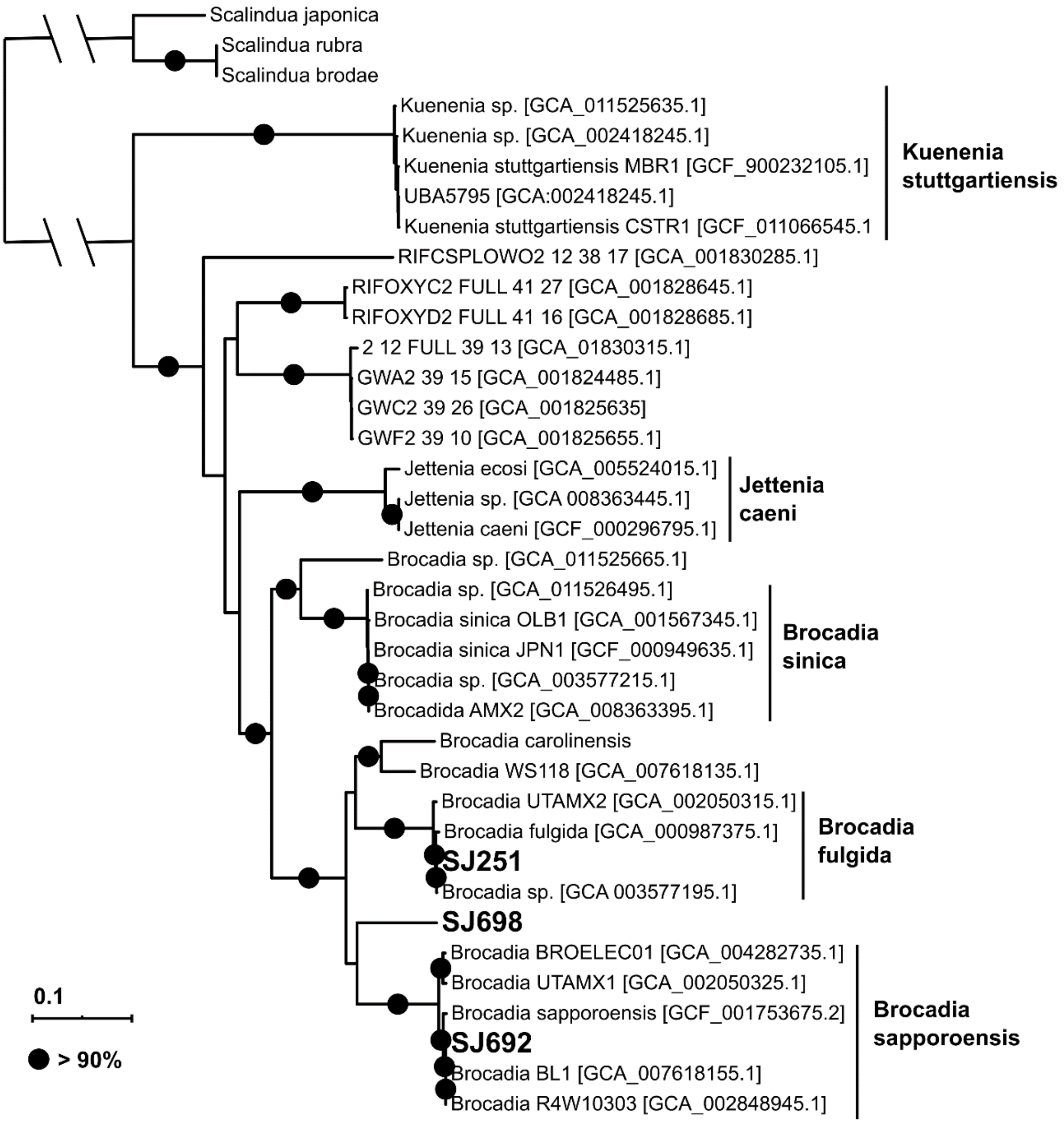

**Figure S8:** Phylogenetic tree for MAGs classified as *Brocadiaaceae* based on 79 single-copy genes. The marine anammox *Scaliduaceae* were used as outgroup. Circles show > 90% ultrafast bootstrap support for a clade.

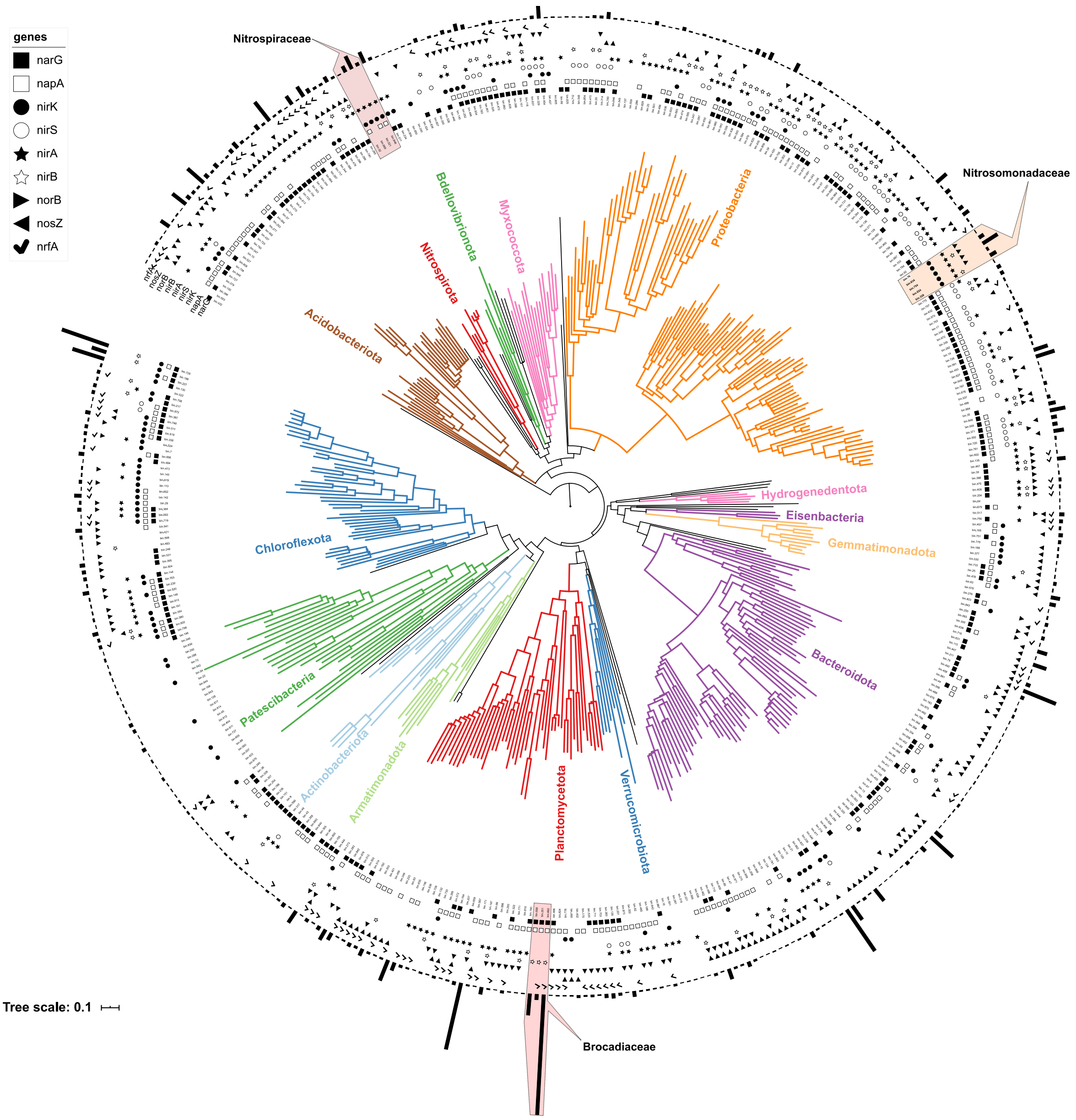

**Figure S9.** Phylogenetic tree of bacterial MAGs using the placement of GTDB-Tk. Bars are proportional to the square transformed average abundance. Symbols represent the presence of genes used for denitrification or nitrite assimilation.

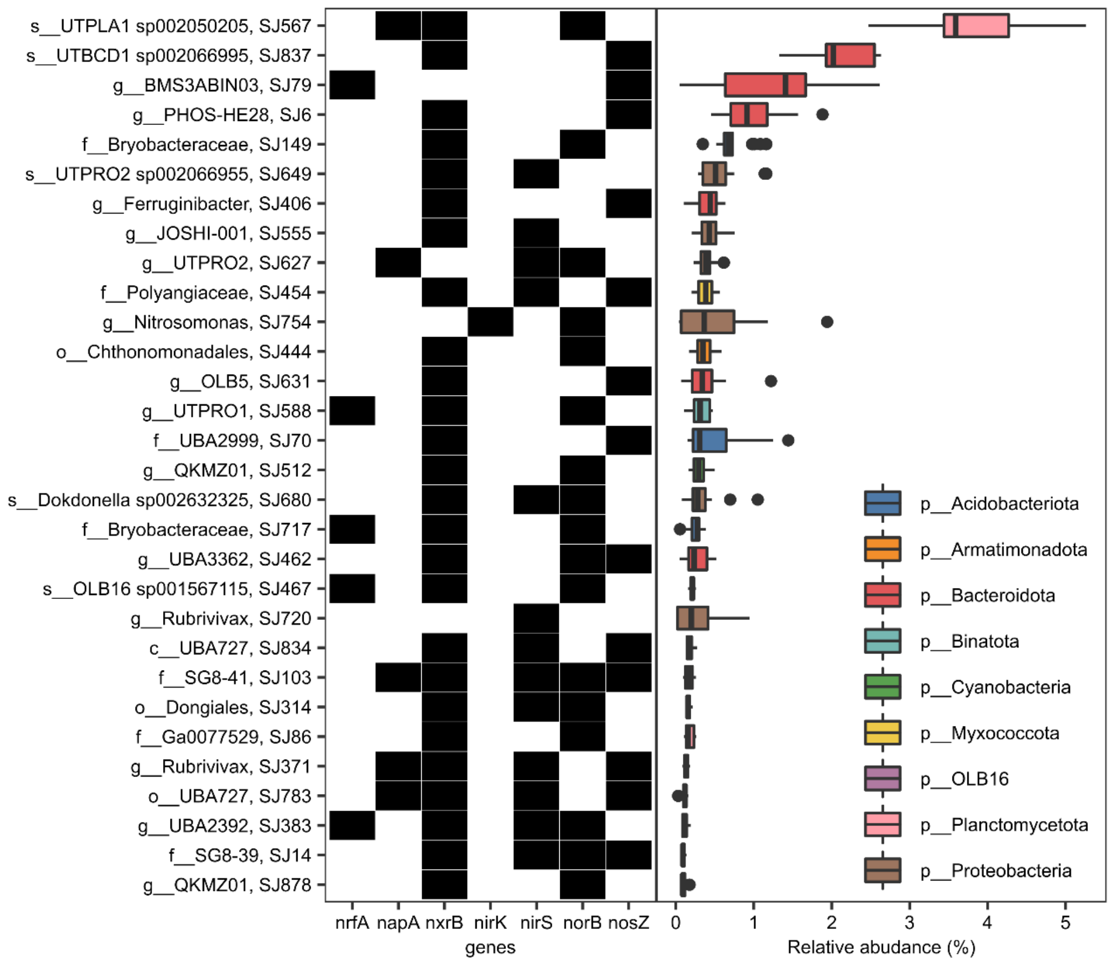

**Figure S10:** Relative abundance of the most abundant denitrifiers, and presence of genes used for denitrification or nitrite assimilation.

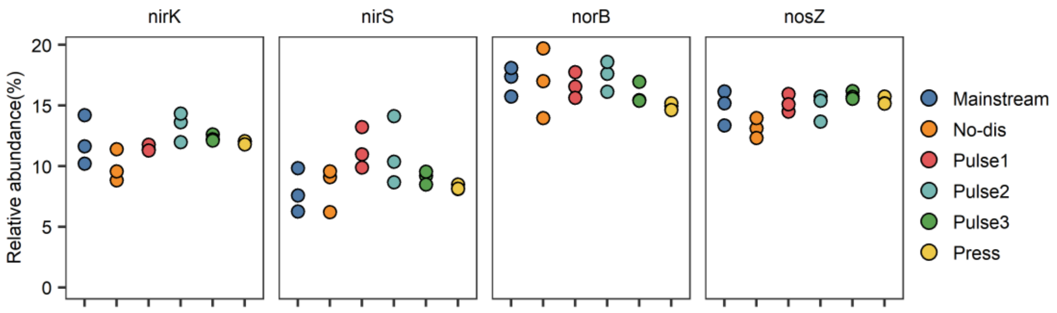

**Figure S11:** Total relative abundance of MAGs with *nirK*, *nirS*, *norB* and *nosZ* genes.

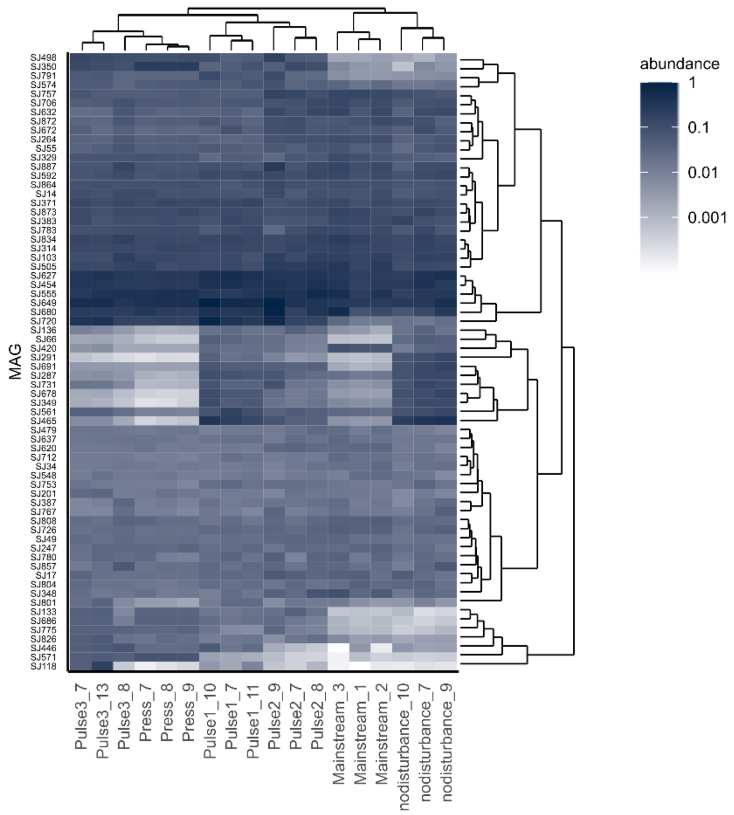

**Figure S12:** Relative abundance of MAGs with the *nirS* genes. Samples and MAGs were clustered by the Bray-Curtis distance of the square transformed data.

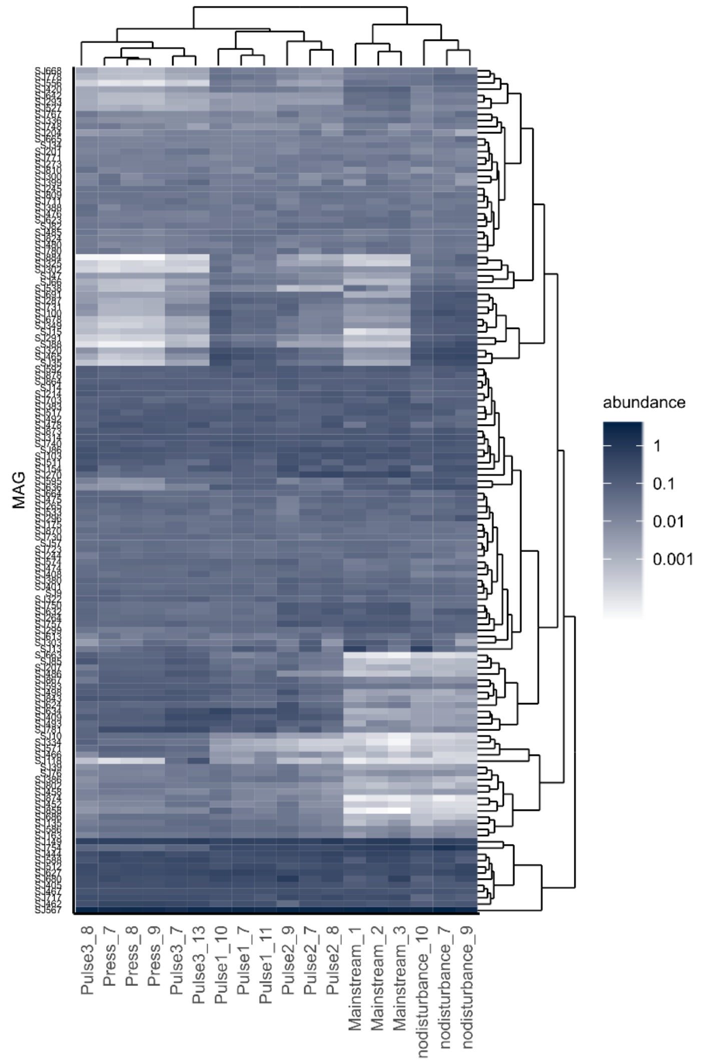

**Figure S13:** Relative abundance of MAGs with a *norB* gene. Samples and MAGs were clustered by the Bray-Curtis distance of the square transformed data.

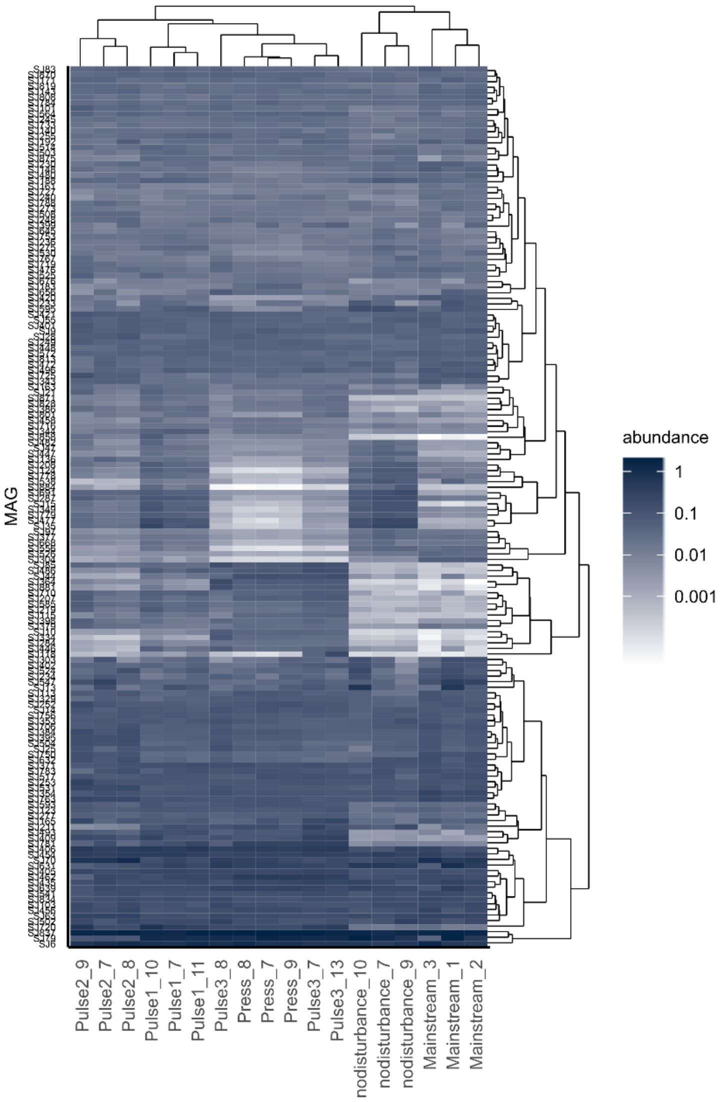

**Figure S14:** Relative abundance of MAGs with a *nosZ* gene. Samples and MAGs were clustered by the Bray-Curtis distance of the square transformed data.

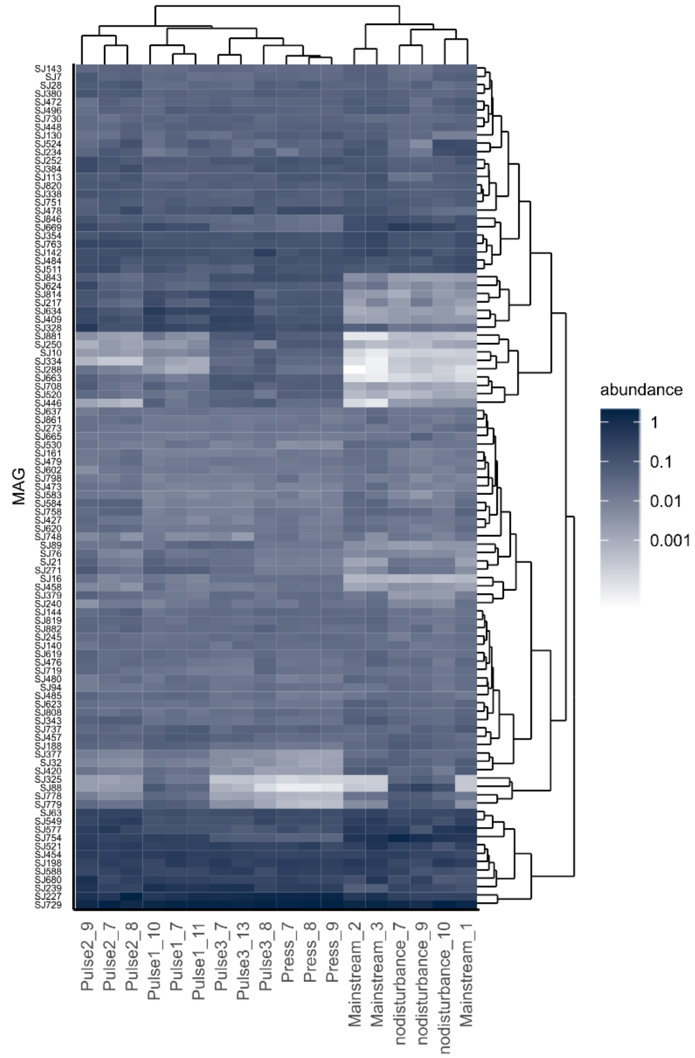

**Figure S15:** Relative abundance of MAGs with a *nirK* gene. Samples and MAGs were clustered by the Bray-Curtis distance of the square transformed data.

### SUPPORTING TABLES

**Table S1:** Conductivity and concentration of ions in sidestream and mainstream wastewater at the Sjölunda WWTP.

| **Sample** | **Date** | **Conductivity** | **$Na^+$** | **$Cl^-$** | **$NH_4^+$** | **$K^+$** | **$PO_4^{3-}$** | **$Mg^{2+}$** | **$Ca^{2+}$** | **$SO_4^{2-}$** |
|---|---|---|---|---|---|---|---|---|---|---|
| | YYMMDD | mS/cm | mM | mM | mM | mM | mM | mM | mM | mM |
| Mainstream | 210117 | 1.23 | 3.99 | 2.92 | 1.06 | 0.39 | 0.63 | 0.79 | 1.86 | 1.05 |
| Mainstream | 210118 | 1.23 | 6.21 | 4.06 | 1.69 | 0.52 | 0.86 | 1.00 | 2.34 | 1.20 |
| Mainstream | 210119 | 1.38 | 7.67 | 4.89 | 1.96 | 0.56 | 0.76 | 0.91 | 2.48 | 1.01 |
| Sidestream | 210118 | 6.97 | 5.74 | 3.79 | 64.6 | 2.69 | 1.43 | 1.98 | 3.70 | 0.50 |
| Sidestream | 210119 | 7.32 | 5.62 | 3.80 | 70.5 | 2.77 | 1.39 | 1.94 | 4.14 | 0.12 |
| Sidestream | 210120 | 7.13 | 6.58 | 4.37 | 68.6 | 2.93 | 1.51 | 2.06 | 3.99 | 0.28 |

**Table S2:** Phylogenetic signal for genes involved in denitrification or nitrite assimilation. The D statistic is an indication of the phylogenetic signal in a binary trait. Values close to or below zero indicate a phylogenetically conserved trait, while values close to one indicate randomness.

| **Gene** | **D** | **N** |
|---|---|---|
| *nirS* | -0.105 | 68 |
| *nirK* | 0.011 | 101 |
| *nosZ* | 0.113 | 158 |
| *nirA* | 0.252 | 172 |
| *nrfA* | 0.266 | 97 |
| *napA* | 0.390 | 230 |
| *norB* | 0.410 | 136 |
| *narG* | 0.494 | 237 |
| *nirB* | 0.518 | 78 |

**Table S3:** Conditions in the MBBRs during the period of investigation

| | Mainstream | Sidestream |
|---|---|---|
| Reactor $NH_4^+$ (mg N $L^{-1}$) | 10 ± 3.0 | 73 ± 24 |
| Reactor $NH_3$ – FA (mg N $L^{-1}$) | 0.04 ± 0.02 | 0.6 ± 0.2 |
| Reactor $NO_2^-$ (mg N $L^{-1}$) | 0.45 ± 0.09 | 4.6 ± 0.7 |
| Reactor $HNO_2$ – FNA (mg N $L^{-1}$) | $9 \pm 2{\cdot}10^{-5}$ | $7 \pm 1{\cdot}10^{-4}$ |
| Reactor $NO_3^-$ (mg N $L^{-1}$) | 3.2 ± 1.1 | 140 ± 28 |
| Temp. (°C) | 15 ± 1.4 | 25 ± 2.2 |
| Dissolved Oxygen (mg $L^{-1}$) | 2.0 ± 0.2 | 1.3 ± 0.7 |
